# Interactions between human milk components and infant polygenic risk predict childhood atopy

**DOI:** 10.64898/2026.08.11.744219

**Authors:** Zhi Yi Fang, Sara A. Stickley, Jihoon Choi, Elizabeth George, Joel Sagman, Amanda M. Zacharias, Amirthagowri Ambalavanan, Charisse Petersen, Bianca Robertson, Chloe Yonemitsu, Kozeta Miliku, Catherine J. Field, Piushkumar J. Mandhane, Elinor Simons, Theo J. Moraes, Michael G. Surette, Lars Bode, Padmaja Subbarao, Stuart E. Turvey, Meghan B. Azad, Qingling Duan

## Abstract

**Background:** Although human milk (HM) confers important health benefits, how bioactive milk components (e.g., microbiota, oligosaccharides, and fatty acids) interact with infant genetics to influence childhood atopy remains poorly understood.

**Objective:** We investigated interactions between infant genomic susceptibility and exposure to maternal human milk components (HMCs) and assessed whether integrating these genetic and milk features improves prediction of childhood atopy.

**Methods:** Leveraging infant genomic and maternal HMC data from the CHILD Cohort Study, we conducted gene-milk interaction analysis using linear regression models that integrated polygenic risk scores (PRS) of nursing infants with multiple HMC types. Gradient-boosting machines (GBMs) were used to evaluate predictive performance of HMCs and infant PRS for childhood atopy.

**Results:** Childhood atopy was associated with interactions between infant genomics (e.g., PRS associated with atopy) and exposure to specific human milk microbes (e.g., *Abiotrophia*, P_Bonf_=0.005, β=0.29), as well as networks of co-occurring HMCs (e.g., a module containing *Bifidobacterium longum*, 2’-fucosyllactose, and eicosapentaenoic acid, P=0.009, β=-12.3). A GBM integrating HMCs and infant PRS achieved the highest predictive performance for childhood atopy with an area under the curve (AUC) of 0.78, outperforming models based on individual HMC types or PRS alone (AUC range: 0.54-0.63).

**Conclusion:** Integration of maternal HMC exposures with infant genomics reveals interaction effects that contribute to prediction of childhood atopy. Understanding how early-life exposures such as HMCs impact the health of children differently depending on their genomic profiles may facilitate the development of personalized intervention strategies to reduce the burden of these health outcomes during childhood.

**Key messages:**

- Interactions between infant polygenic risk and exposure to human milk components are associated with childhood atopy.
- Networks of co-occurring human milk microbiota, oligosaccharides, and fatty acids may influence childhood atopy, with effects varying by infant genomic susceptibility.
- Integration of human milk components with infant genomics improves prediction of childhood atopy compared with individual milk components or genomics alone.

**Capsule Summary:** This study demonstrates that interactions between infant polygenic risk and maternal milk components improve prediction of childhood atopy, highlighting opportunities for personalized early-life prevention strategies.

## Introduction

Human milk (HM) is a biological fluid enriched with bioactive components that support infant immune and physiological development.^1^ Among these, the human milk microbiota (HMM) has been implicated in shaping infant immune maturation, which may influence susceptibility to allergic disease later in life.^2^ Our team recently reported associations between specific HMM traits, such as variation in the abundance of *Granulicatella sp.*, and childhood sensitization at ages 1-5 years.^3^ Emerging evidence further suggests that exposure to HM traits may influence atopy risk differently among milk-fed children depending on their genomic profiles.^4^ Further investigation is needed, however, into the additive and interaction effects of infant genomics with exposure to HMM traits on risk of childhood atopy.

In addition to microbiota, HM contains other bioactive components, such as fatty acids (HMFAs) and oligosaccharides (HMOs), which may modify the composition of HMM.^5^ For example, increased abundance of the HMFA omega-3 (n-3) polyunsaturated fatty acids (PUFAs) has been associated with reduced *Bifidobacterium* abundance in HM.^6^ In addition, a recent study reported that 19 of the most abundant HMOs are metabolized by over 100 bacterial species in the gut.^7^ While most studies have focused on dietary fatty acids or HMOs associated with infant gut microbiota,^7,8^ few have investigated the influence of HMFAs or HMOs on individual microbes or communities within the HMM.^5,6,9^ Moreover, although HMOs, HMFAs, and HMM have each been associated with childhood health outcomes,^3,10,11^ their combined (additive and multiplicative) effects on childhood atopy remain unknown.

We hypothesized that interactions between infant genomic susceptibility and exposure to HM components (HMCs) influence childhood atopy. First, we determined which HMCs may be linked to childhood atopy by assessing gene-milk interaction effects on childhood atopy using infant polygenic risk scores (PRS) and variable HMCs from lactating mothers. We then assessed whether HMM associated with atopy was influenced by HMOs and HMFAs. Finally, unsupervised and supervised machine learning approaches were applied to investigate the combined impact of HMCs and infant PRS on childhood atopy. Our nuanced study elucidates the complex interactions among the components of the mother-milk-infant triad.^12^ By integrating infant genomics with multiple types of HMCs, we demonstrate that their interactions modulate atopic health and improve prediction of childhood atopy. Understanding how HMCs influence childhood atopy across variable PRS levels could inform personalized interventions during early life.

## Methods

### CHILD Cohort Study Participants

This study leveraged data from the CHILD Cohort Study, a prospective birth cohort of over 3,000 families recruited across four sites (Vancouver, Edmonton, Winnipeg, and Toronto) (**Figure 1A**).^13^ Women (N=3,624) with singleton pregnancies were recruited during gestation, and infants were eligible if born at ≥35 weeks of gestation and with a birth weight ≥2,500 g (**Figure 1B**).^13^ The current study included 885 mothers with HMM data and 689 of their children who were fed HM for at least 6 months, had genomic data, and health information from ages 1 to 5 years (**Figure E1**).^13^ Study protocols were approved by the Human Research Ethics Boards at Queen’s University, McMaster University, the Universities of Manitoba, Alberta and British Columbia, and the Hospital for Sick Children.

**Figure 1.**
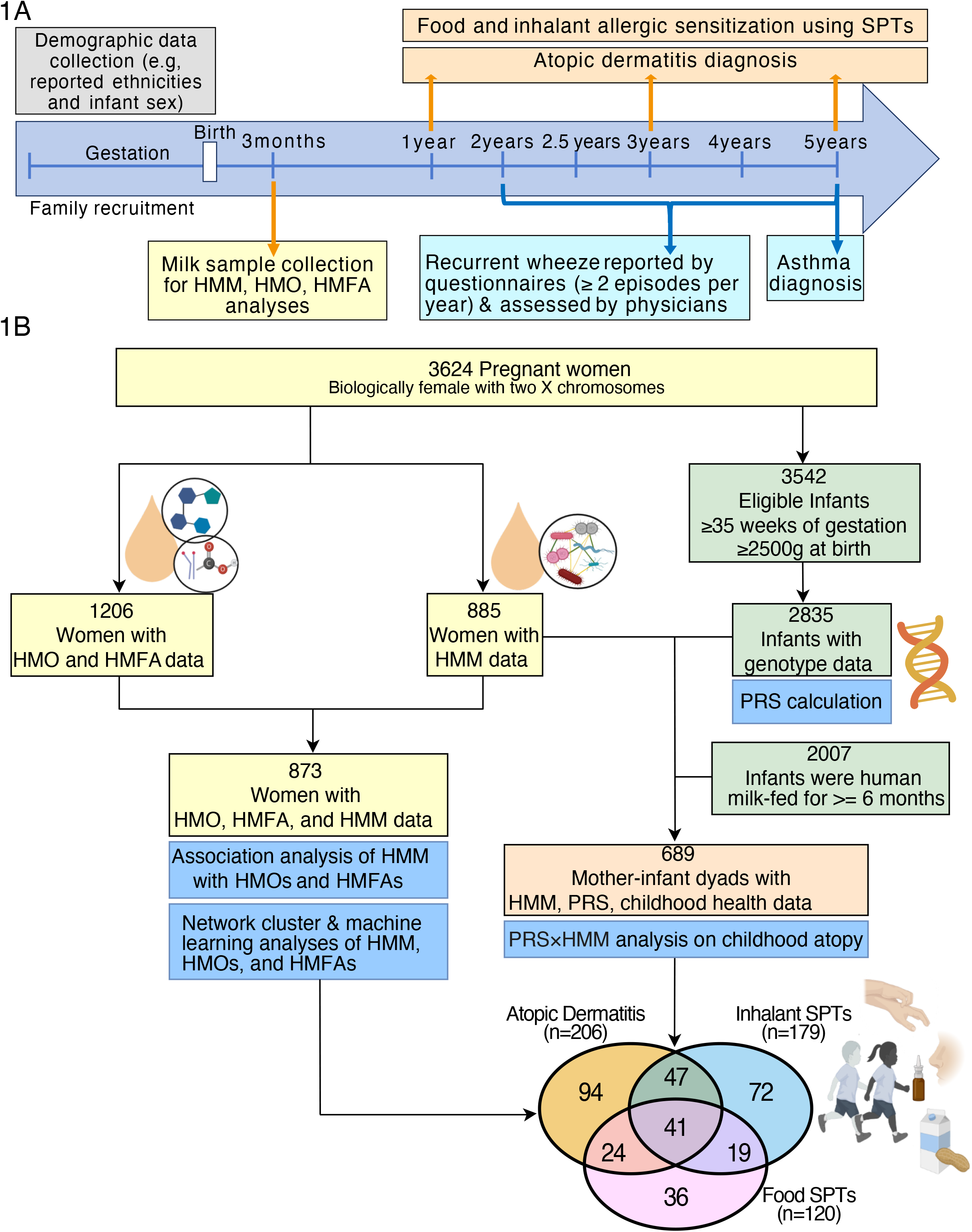
Study Overview. (**A**) Enrollment and data collection timeline for the CHILD Cohort Study, spanning from gestation to age five years. (**B**) A total of 885 mothers had HMM profiles available from 16S rRNA sequencing. Of these, 689 had children with health information from ages 1-5 years who were fed HM for at least 6 months.

### Childhood atopy and related outcomes

Childhood atopy was defined as physician-diagnosed atopic dermatitis (AD)^14,15^ and/or sensitization to food or inhalant allergens, as determined by positive SPTs at ages 1, 3, and 5 years (**Figure 1A**).^16^ Given that the majority of childhood atopy co-occurs with asthma,^17^ we derived a secondary outcome variable to include any atopy at ages 1-5 years, asthma at age 5 years, and recurrent wheeze between ages 2-5 years.^13^ The “control” group consisted of children (n=214) without any of these conditions at any assessed age, and the case group included children (n=385) with ≥1 condition (**Table E1**). As previously described, recurrent wheeze was identified in children who experienced ≥2 episodes of wheezing within one year (**Figure 1A**). See **Supplementary Methods** for details.

### HM collection and quantification of HMCs

HM samples were collected in two batches and processed similarly as previously described.^18,19^ Each mother provided one milk sample from multiple feeds collected over 24-hours at 3-6 months postpartum. HMM profiling of 885 HM samples has been described previously (**Figure 1**).^20,21^ Microbial composition was characterized using 16S rRNA gene sequencing targeting the V4 hypervariable region.^22,23^ Sequence preprocessing^22,24,25^, quality control (QC),^26,27^ and taxonomic annotation^28^ were performed following established pipelines.^3,21^ ASVs and PICRUSt2^29^-predicted metabolic pathways were filtered for >5% prevalence,^3^ resulting in 179 ASVs and 378 pathways (**Figure E1**). Abundances were centered-log-ratio (CLR) transformed after zero-value imputation (**Figure E1**).^3,30^ Alpha-diversity (i.e., Shannon) and six co-abundance HMM network modules derived from 179 ASVs were previously imputed to capture community-level variation in HMM (**Figure E1**).^3^ See **Supplementary Methods** for details.

The 19 and 28 most abundant HMOs and HMFAs, respectively, were quantified from approximately 1,200 mothers using high-performance liquid chromatography^18^ and gas-liquid chromatography,^31^ respectively. Additionally, total HMO concentrations were calculated as the sum of all 19 measured HMOs, while HMO-bound fucose- and sialic acid concentrations were calculated by summing fucose and sialic acid residues, respectively.^31^ Summary measures and ratios of HMFAs were previously calculated.^31^ Rank-based inverse normal transformation was applied to HMO and HMFA data using the RNOmni R package.^32^ See **Supplementary Methods** and **Tables E2** and **E3** for details.

### Infant genomics and PRS calculation

Genomic DNA from cord blood of 2,967 infants was genotyped using the Illumina HumanCoreExome BeadChip.^10^ Following QC using PLINK,^10,33,34^ genotypes from 2,835 infants were imputed to the Haplotype Reference Consortium r1.1 panel on the Michigan server (**Figure 1B**).^35^ PRS for atopic outcomes in the CHILD dataset were calculated using Polygenic Score Catalog Calculator (pgsc_calc v2.0.0)^36^ with weights derived from the genome-wide association studies of allergic diseases (PGS ID: PGS001285) (**Figure 1B**).^37^ See **Supplementary Methods** for details.

### Statistical analysis

#### Infant PRS and maternal HMM interaction analyses on childhood atopy

Our team has previously investigated the main associations between HMM exposure and childhood atopy (**Tables 1 and E4**).^3^ In this study, gene-milk interaction analyses on childhood atopy at ages 1-5 years were assessed using binomial generalized linear models (GLMs) with a multiplicative interaction term between infant PRS and HMM (PRS×HMM) (**Figure 1B**). Models were adjusted for study center, milk batch, time from milk sample collection to processing, infant sex, antibiotic exposure during the first year of life, and parent-reported infant ethnicity. To account for correlations among HMM features, 85 and 98 independent features were previously estimated for 179 ASVs and 378 metabolic pathways using matSpDlite,^3,38^ and Bonferroni correction was applied accordingly (i.e., α = 0.05/85 for ASVs and α = 0.05/98 for metabolic pathways). HMM traits or modules with significant main^3^ or interaction effects were carried forward for downstream analyses (**Table 1**). The interaction plots were generated using the R package sjPlot (v.2.9.0; https://CRAN.R-project.org/package=sjPlot).

**Table 1.**
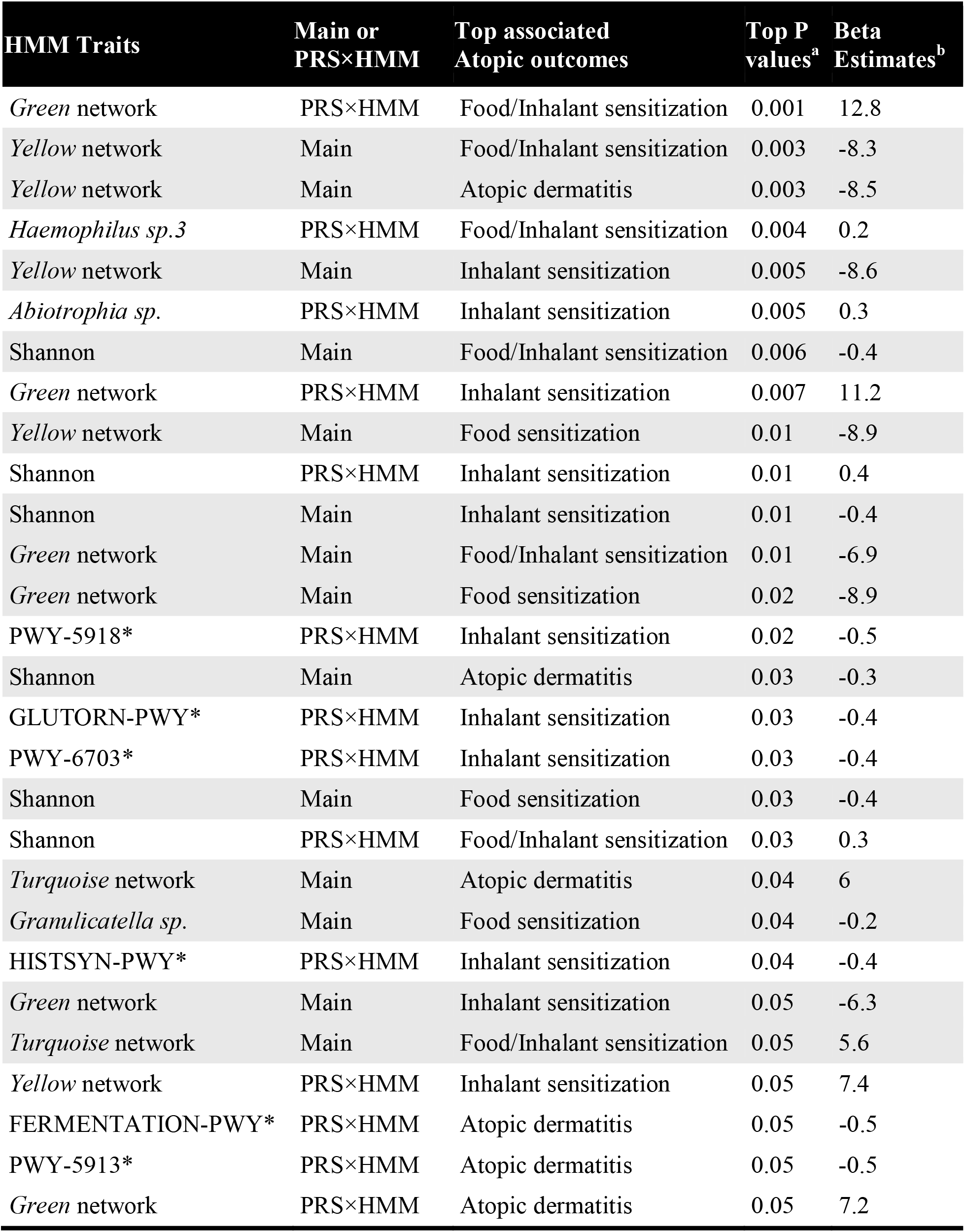
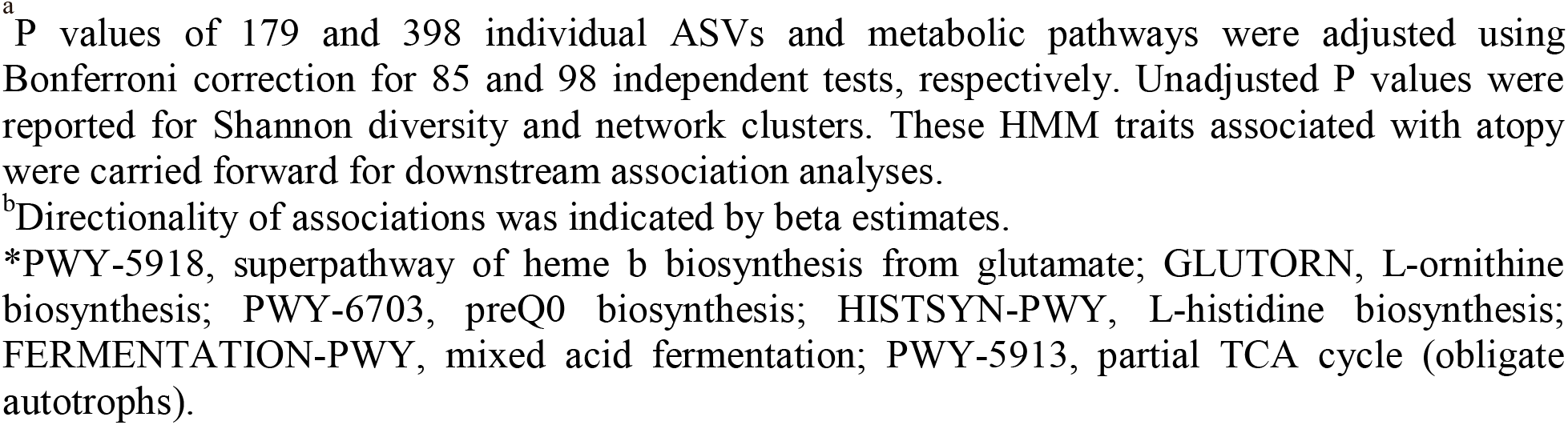
Main and interaction effects between infant PRS and maternal HMM associated with childhood atopy.

| HMM Traits | Main or PRS×HMM | Top associated Atopic outcomes | Top P values<sup>a</sup> | Beta Estimates<sup>b</sup> |
| --- | --- | --- | --- | --- |
| *Green network* | PRS×HMM | Food/Inhalant sensitization | 0.001 | 12.8 |
| *Yellow network* | Main | Food/Inhalant sensitization | 0.003 | -8.3 |
| *Yellow network* | Main | Atopic dermatitis | 0.003 | -8.5 |
| *Haemophilus sp.3* | PRS×HMM | Food/Inhalant sensitization | 0.004 | 0.2 |
| *Yellow network* | Main | Inhalant sensitization | 0.005 | -8.6 |
| *Abiotrophia sp.* | PRS×HMM | Inhalant sensitization | 0.005 | 0.3 |
| Shannon | Main | Food/Inhalant sensitization | 0.006 | -0.4 |
| *Green network* | PRS×HMM | Inhalant sensitization | 0.007 | 11.2 |
| *Yellow network* | Main | Food sensitization | 0.01 | -8.9 |
| Shannon | PRS×HMM | Inhalant sensitization | 0.01 | 0.4 |
| Shannon | Main | Inhalant sensitization | 0.01 | -0.4 |
| *Green network* | Main | Food/Inhalant sensitization | 0.01 | -6.9 |
| *Green network* | Main | Food sensitization | 0.02 | -8.9 |
| PWY-5918* | PRS×HMM | Inhalant sensitization | 0.02 | -0.5 |
| Shannon | Main | Atopic dermatitis | 0.03 | -0.3 |
| GLUTORN-PWY* | PRS×HMM | Inhalant sensitization | 0.03 | -0.4 |
| PWY-6703* | PRS×HMM | Inhalant sensitization | 0.03 | -0.4 |
| Shannon | Main | Food sensitization | 0.03 | -0.4 |
| Shannon | PRS×HMM | Food/Inhalant sensitization | 0.03 | 0.3 |
| *Turquoise network* | Main | Atopic dermatitis | 0.04 | 6 |
| *Granulicatella sp.* | Main | Food sensitization | 0.04 | -0.2 |
| HISTSYN-PWY* | PRS×HMM | Inhalant sensitization | 0.04 | -0.4 |
| *Green network* | Main | Inhalant sensitization | 0.05 | -6.3 |
| *Turquoise network* | Main | Food/Inhalant sensitization | 0.05 | 5.6 |
| *Yellow network* | PRS×HMM | Inhalant sensitization | 0.05 | 7.4 |
| FERMENTATION-PWY* | PRS×HMM | Atopic dermatitis | 0.05 | -0.5 |
| PWY-5913* | PRS×HMM | Atopic dermatitis | 0.05 | -0.5 |
| *Green network* | PRS×HMM | Atopic dermatitis | 0.05 | 7.2 |

<sup>a</sup> P values of 179 and 398 individual ASVs and metabolic pathways were adjusted using Bonferroni correction for 85 and 98 independent tests, respectively. Unadjusted P values were reported for Shannon diversity and network clusters. These HMM traits associated with atopy were carried forward for downstream association analyses.
<sup>b</sup> Directionality of associations was indicated by beta estimates.
\*PWY-5918, superpathway of heme b biosynthesis from glutamate; GLUTORN, L-ornithine biosynthesis; PWY-6703, preQ0 biosynthesis; HISTSYN-PWY, L-histidine biosynthesis; FERMENTATION-PWY, mixed acid fermentation; PWY-5913, partial TCA cycle (obligate autotrophs).

### Association analyses of HMOs and HMFAs with HMM

Associations between HMOs or HMFAs and 13 HMM traits implicated in atopy were assessed using Gaussian GLMs (**Figure 1B**), adjusting for study center, milk batch, time from milk sample collection to processing, and maternal self-reported ethnicity. To account for multiple tests, Bonferroni correction was applied based on the six HMO and three HMFA clusters estimated for the 19 and 28 HMOs and HMFAs, respectively (**Figure E2**).

### Multi-component network analysis

We conducted multi-component network analyses to investigate co-abundance patterns among 179 ASVs, 19 HMOs, and 28 HMFAs using a ‘signed’ network approach in the weighted gene co-expression network analysis R package (v.1.69.81).^39^ Prior to analysis, all HM features were scaled to a common range (–2 to +2) to ensure equal weighting across data types. The resulting network modules were visualized using Cytoscape^40^ and summarized by module eigenvalues derived from principal component (PC) analysis. These module eigenvalues, corresponding to the first PC of each module, were used to summarize the co-occurrence pattern of ASVs, HMOs, and HMFAs within each module.^39^ GLMs were then applied to assess both the main effects of exposure to multi-component network HM modules and interactions with PRS (PRS×HMC) on childhood atopy at ages 1-5 years (**Figure 1**). Models were adjusted for study center, milk batch, time from milk sample collection to processing, infant sex, antibiotic exposure during the first year of life, and infant ethnicity reported by parents using questionnaires. See **Supplementary Methods** and **Figure E3** for details.

### Machine Learning of integrated milk components and PRS for childhood atopy

Gradient-boosting machines (GBMs) were implemented using the caret R package (v.6.0.94)^41^ to evaluate the performance of machine learning models integrating ASVs, HMFAs, and HMOs with infant PRS in classifying cases and controls among infants fed HM for at least 6 months. We compared models trained on individual HMC types or PRS alone with the model incorporating all HMCs and PRS.^41^ Prior to model training, all features were standardized (mean = 0, SD = 1), and feature selection was based on significant main and interaction effects identified in GLMs (**Table E5**). GBMs were trained using a 70:30 train-test split with five repeats of 3-fold cross-validation.^41^ Model performance was evaluated using the area under the curve (AUC) calculated using the ROCR R package (v.1.0.11),^42^ and variable importance was assessed using *varImp()* in the caret R package (v.6.0.94)^41^. See **Supplementary Methods** and **Table E5** for details.

## Results

### CHILD Study participants

This study included 885 mothers from the CHILD Cohort Study, a longitudinal birth cohort of >3,000 families across four Canadian sites. Hand-expressed HM samples collected at 3-6 months postpartum were used for targeted 16S rRNA sequencing to profile HMM (**Figure 1**). Of these mothers, 873 had HMFA and HMO data, and 689 of their children, who were fed HM for at least six months, had genomic data and clinical follow-up from ages 1 to 5 years (**Figure E1**). Childhood atopy, defined as physician-diagnosed AD and/or positive SPTs, was assessed at ages 1, 3, and 5 years (**Figure 1**). We curated a secondary outcome for machine learning analyses to capture broader allergy-related phenotypes: presence of any atopy between ages 1-5 years, asthma at age 5 years, or recurrent wheeze between ages 2-5 years (385 cases vs. 214 controls) (**Table E1**).

### Interactions between infant PRS and maternal HMM were associated with childhood atopy

Binomial GLMs identified significant interactions between infant PRS and HMM traits (i.e., two ASVs and six metabolic pathways) associated with childhood atopy (**Figure 2, Tables 1 and E4**). In these GLMs, the interaction β coefficient represents how the association between exposure to an HMM trait and childhood atopy changes with increasing PRS: among infants with high PRS, a positive β indicates that exposure to increased HMM abundance is associated with higher atopy prevalence, whereas a negative β indicates that exposure to increased HMM abundance is associated with reduced atopy prevalence. For example, exposure to increased abundance of *Haemophilus sp. 3* and *Abiotrophia sp.* in HM was associated with increased prevalence of allergic sensitization (i.e., positive SPTs) among HM-fed infants with high PRS for atopy (P_Bonf_=0.004, β=0.16; **Figure 2A**). Among infants with low to moderate PRS, the same exposure was associated with reduced prevalence of allergic sensitization (**Figure 2A**). We identified six metabolic pathways with protective associations against atopy among infants with high PRS (**Figure 2A and B**): L-histidine biosynthesis (HISTSYN-PWY; P_Bonf_ = 0.04, β = -0.37), preQ0 biosynthesis (PWY-6703; P_Bonf_=0.03, β=-0.4), L-ornithine biosynthesis (GLUTORN-PWY; P_Bonf_=0.03, β=-0.36), superpathway of heme biosynthesis from glutamate (PWY-5918; P_Bonf_=0.02, β=-0.51), partial TCA cycle (obligate autotrophs) (PWY-5913; P_Bonf_=0.05, β=-0.5), and mixed acid fermentation (FERMENTATION-PWY; P_Bonf_=0.05, β=-0.53). Among infants with low to moderate PRS, these same exposures were associated with increased prevalence of childhood allergic sensitization (**Figure 2A and B**).

**Figure 2.**
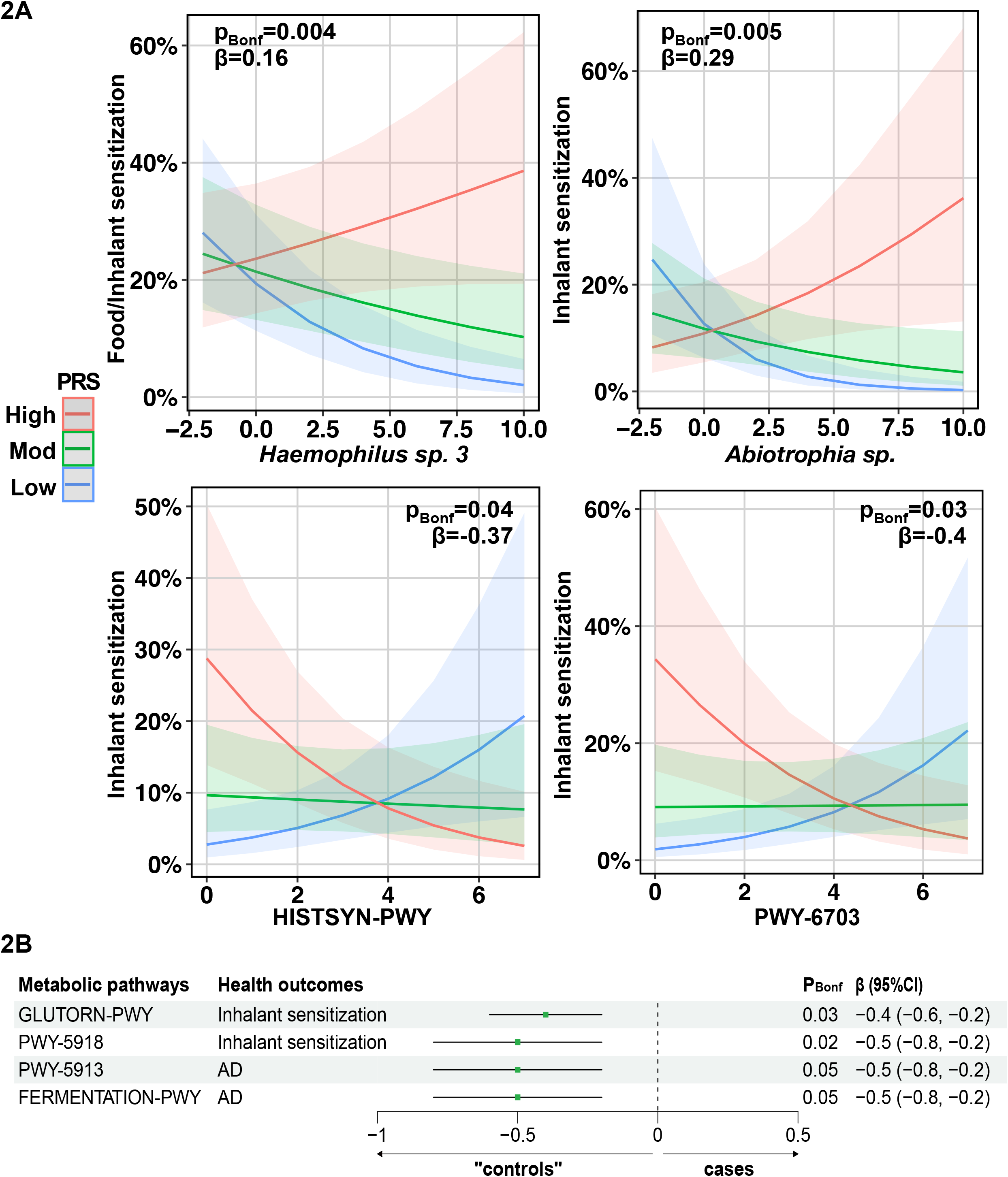
Interactions between infant PRS and HMM traits were associated with childhood atopy. CLR-transformed abundances of individual ASVs (**A**) and MetaCyc metabolic pathways (**A** and **B**) were associated with childhood atopy depending on infant PRS. P_Bonf_ represents Bonferroni-adjusted P values. The interaction beta (β) coefficients indicate how the association between exposure to HMM traits and childhood atopy changes with increasing PRS. PRS were categorized into low (<-1 SD), moderate (Mod; -1 to +1SD), and high (>+1 SD).

At the community level, increased Shannon diversity (P=0.01, β=0.39; **Figure 3A**, **Table E4**) and abundance of co-occurring milk taxa in the *yellow* and *green* (P=0.007, β=11.2; **Figure 3A and B**, **Table E4**) modules were associated with increased prevalence of sensitization to inhalant allergens among HM-fed infants with high PRS for atopy. Our interaction results demonstrate that exposure to HMM traits alone may not have a main effect on childhood atopy, but it may exert different effects on atopy depending on the infants’ individual PRS. These results for HMM contrast with our analysis of breastfeeding duration (6 months), which showed no significant interaction with infant PRS on atopy risk.

**Figure 3.**
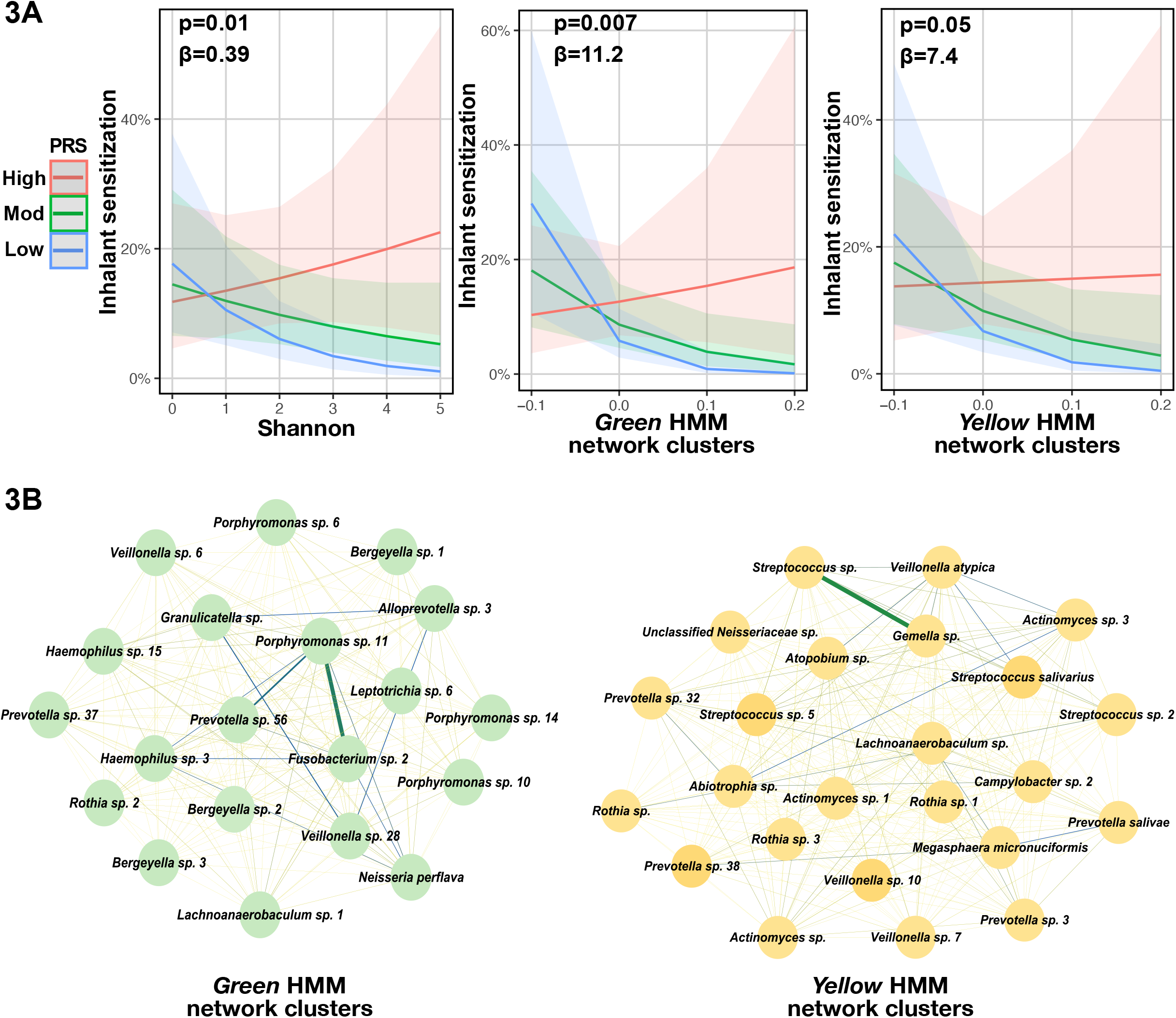
Interactions between infant PRS and HMM community were associated with childhood atopy. Shannon diversity and HMM network modules (**A**) were associated with childhood atopy depending on infant PRS. P represents unadjusted P values. The interaction beta (β) coefficients indicate how the association between exposure to HMM traits and childhood atopy changes with increasing PRS. PRS were categorized into low (<-1 SD), moderate (Mod; -1 to +1 SD), and high (>+1 SD). (**B**) shows that the HMM *green* and *yellow* network modules contained 20 and 24 ASVs, respectively. Edges (lines) between ASVs (nodes) vary in width and colour: thicker and green lines represent stronger associations and thinner and yellow lines represent weaker associations.

### HMFAs and HMOs were associated with atopy-implicated HMM traits

Our Gaussian GLMs showed that mothers with decreased concentrations of omega-3 and omega-6 fatty acids, such as eicosatetraenoic acid (20:4n3), eicosapentaenoic acid (20:5n3), and arachidonic acid (20:4n6) had increased abundance of *Abiotrophia sp.* (P_Bonf_=5.8e-4, β=-0.25) and *Granulicatella sp.* (P_Bonf_=7.7e-3, β=-0.23) in their milk (**Figure 4A, Table E6**). Increased concentrations of 20:4n3 and 20:5n3 were also associated with increased abundance of HISTSYN-PWY (P_Bonf_ = 5.1e-4, β = 0.14). Moreover, mothers with increased total omega-6 to omega-3 polyunsaturated fatty acid (n-6/n-3 PUFA) ratio had increased abundances of *Abiotrophia sp.* (P=0.03, β=0.15) and ASVs within the *green* and *yellow* modules (P=0.03, β=0.003). Similarly, increased total arachidonic acid to docosahexaenoic acid (ARA/DHA) ratio was also associated with increased abundance of *Abiotrophia sp.* (P=0.04, β=0.15).

**Figure 4.**
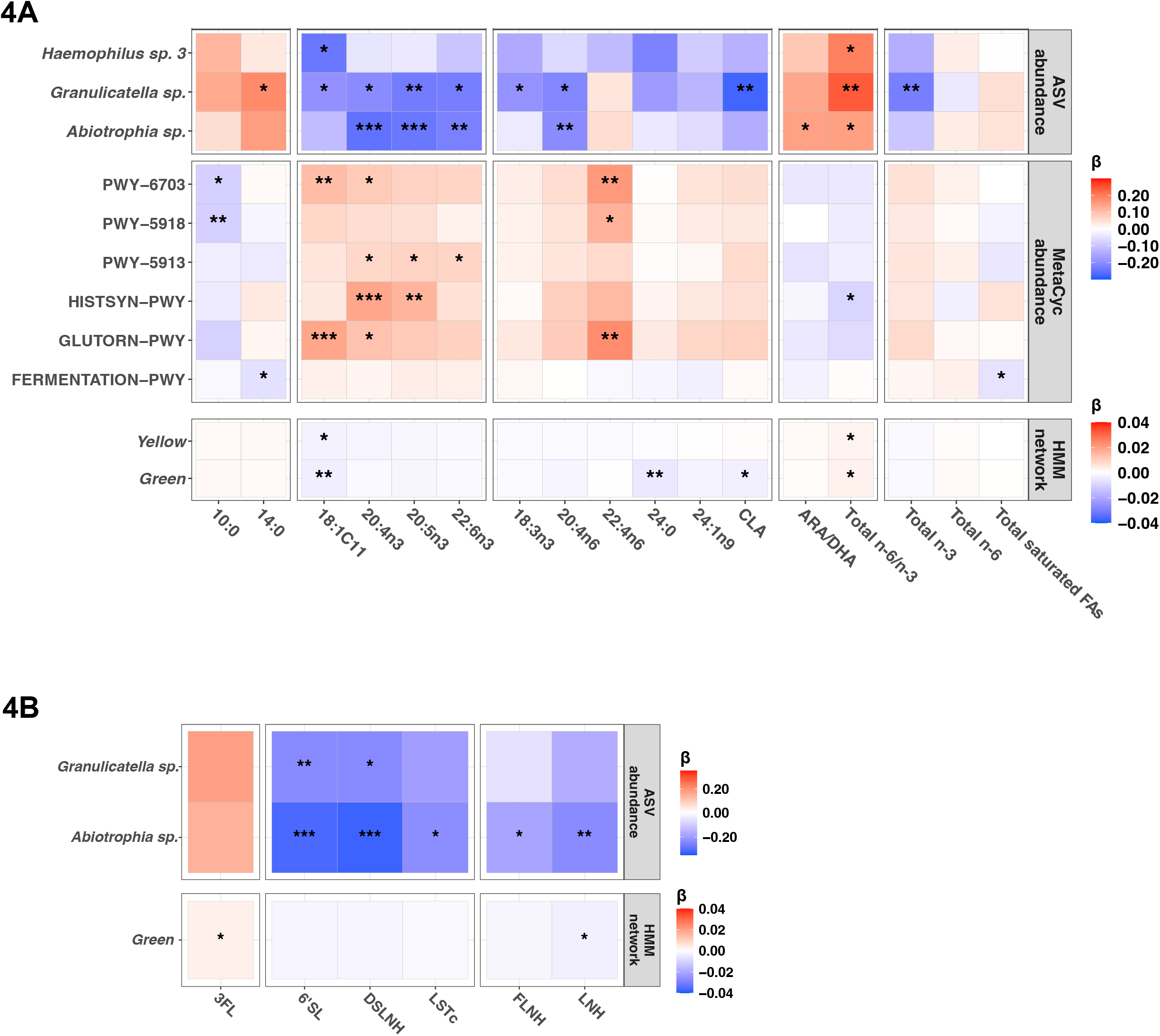
HMOs and HMFAs were associated with atopy-implicated HMM. CLR-transformed abundances of individual ASVs and metabolic pathways, as well as the ASVs within the *green* HMM cluster, were associated with HMFAs (**A**) and HMOs (**B**). For associations with individual HMOs or HMFAs, *P <0.05; **P <0.01; ***P <0.001 represent Bonferroni-adjusted P values based on three clusters of correlated HMFAs and six clusters of correlated HMOs, as estimated in (**A** and **B**). For associations with sums or ratios of HMOs or HMFAs, *P <0.05; ** P <0.01; ***P <0.001 represent unadjusted P values in (**A** and **B**). A separate beta (β) estimate legend was generated for ASV/MetaCyc abundance (top) and HMM network (bottom) in each panel. Positive β (red) indicates that increased HMO or HMFA concentrations are associated with increased abundance of HMM traits. Negative β (blue) indicates that decreased HMO or HMFA concentrations are associated with increased abundance of HMM traits. Colour intensity represents the magnitude of β. 10:0, capric acid; 14:0, myristic acid; 18:1c11, cis-vaccenic acid; 20:4n3, eicosatetraenoic acid; 20:5n3, eicosapentaenoic acid; 22:6n3, docosahexaenoic acid (DHA); 18:3n3, α-linolenic acid; 20:4n6, arachidonic acid (ARA); 22:4n6, docosatetraenoic acid; 24:0, lignoceric acid; 24:1n9, nervonic acid; CLA, conjugated linoleic acid; ARA/DHA, ARA to DHA ratio; Total n-6/n-3, omega-6 to omega-3 polyunsaturated fatty acid (PUFA) ratio; Total n-3, total omega-3 PUFAs; Total n-6, total omega-6 PUFAs; Total saturated FAs, total saturated fatty acids. 3-fucosyllactose, 3FL; 6′-sialyllactose, 6′SL; disialyllacto-N-hexaose, DSLNH; LS-tetrasaccharide c, LSTc; fucosyllacto-N-hexaose, FLNH; lacto-N-hexaose, LNH.

In addition to HMFAs, we determined that decreased concentrations of the HMO lacto-N-hexaose (LNH) were associated with increased abundances of *Abiotrophia sp.* (P_Bonf_=0.002, β=-0.24) and ASVs within the *green* network module (P_Bonf_=0.03, β=-0.003) (**Figure 4B**, **Table E7**). Increased abundance of *Abiotrophia sp.* was also associated with decreased concentrations of three sialylated HMOs: 6’-sialyllactose (6′SL), disialyllacto-N-hexaose (DSLNH), and LS-tetrasaccharide c (LSTc) (P_Bonf_=5.3e-6, β=-0.33). Thus, our results demonstrate that HMFAs and HMOs are associated with atopy-implicated HMM.

### Interactions between infant PRS and multi-component network modules were associated with childhood atopy

To investigate co-occurrence patterns among HMCs, we clustered 19 HMOs, 28 HMFAs, and 179 ASVs into seven network modules (**Figures 5 and E3**, **Table E8**). Three (i.e., *multi-red*, *multi-blue*, and *multi-brown*) of the seven modules were made up of more than one HMC type. These three modules were further assessed for main and interaction effects on childhood atopy. We observed a strong association between decreased prevalence of childhood sensitization to both food and inhalant allergens and exposure to increased HMO and ASV abundances (e.g., *Abiotrophia sp.*, *Veillonella atypica*, and 3-fucosyllactose (3FL)) within the *multi-blue* module (P=0.006, β=-10.6) (**Figure 6A**) among infants with low or moderate PRS (P=0.001, β=11.8) (**Figure 6B and C**). Our PRS interaction analyses showed that this module was not associated with atopy prevalence among those with high PRS, suggesting that exposure to these HMCs may not modulate atopic risk in children with a stronger genetic predisposition (**Figure 6B and C**). In addition to the *multi-blue* module, our interaction analysis also determined that the exposure to increased HMO, HMFA, and ASV abundances within the *multi-brown* module (e.g., 2′-fucosyllactose (2′FL), 6′SL, eicosapentaenoic acid (20:5n3), adrenic acid (22:4n6), *Bifidobacterium longum 1*) was associated with reduced prevalence of food sensitization at ages 1-5 among infants with high PRS (P=0.009, β=-12.3; **Figure 6B**). Overall, our findings indicate that infant PRS may modify the associations of network modules composed of certain HMOs, HMFAs, and ASVs with childhood atopy.

**Figure 5.**
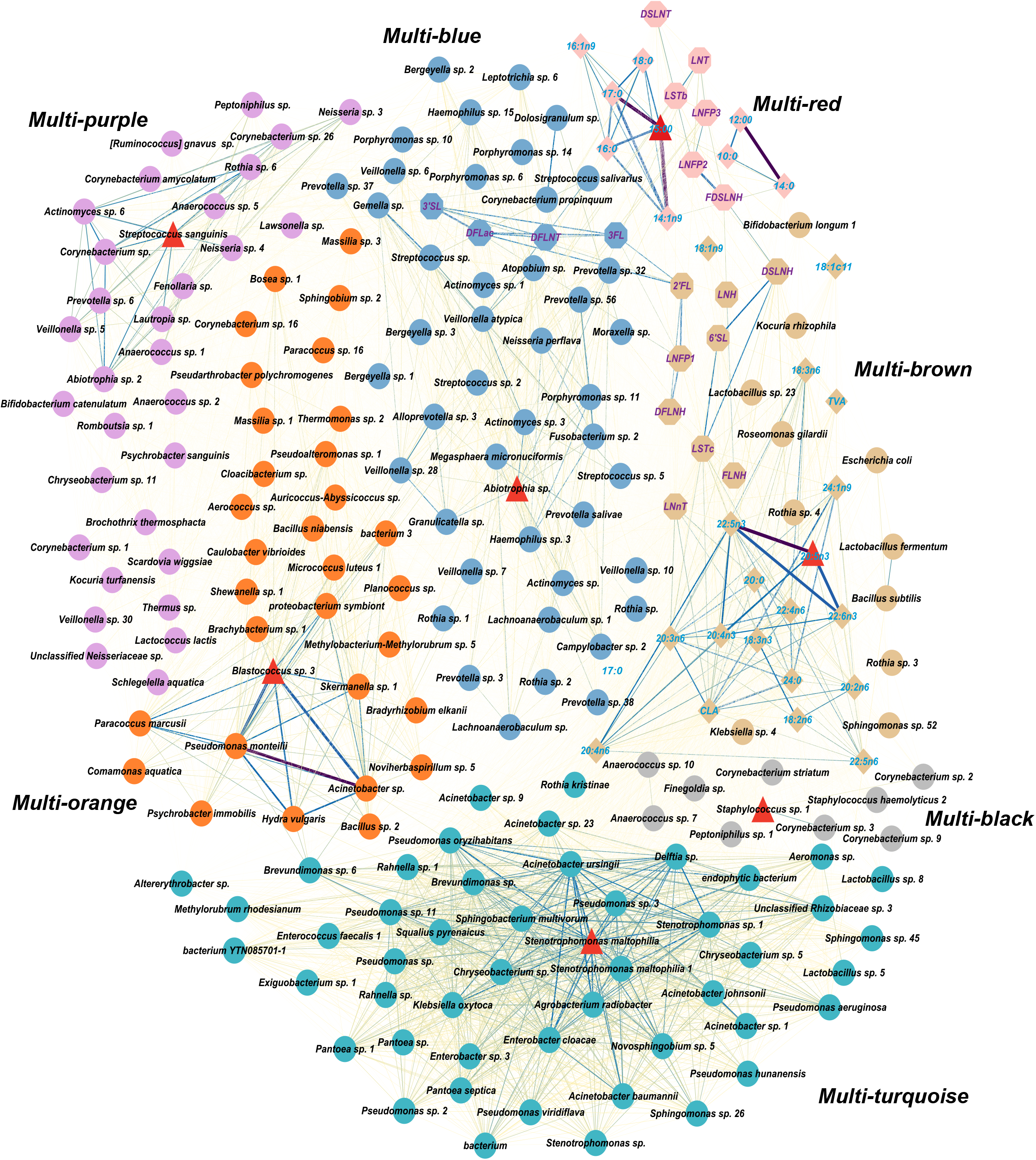
Seven multi-component network modules of correlated HMOs, HMFAs, and ASVs. A total of 19 HMOs (octagons), 28 HMFAs (diamonds), and 179 HM ASVs (circles) were clustered into seven network modules, each indicated by a different colour. Red triangles denote hub milk features with the highest intramodular connectivity. Edges (lines) between the human milk components (nodes) vary in width and colour: thicker and purple lines represent stronger associations and thinner and yellow lines represent weaker associations. The number of co-occurring human milk components (n) within each module is labelled.

**Figure 6.**
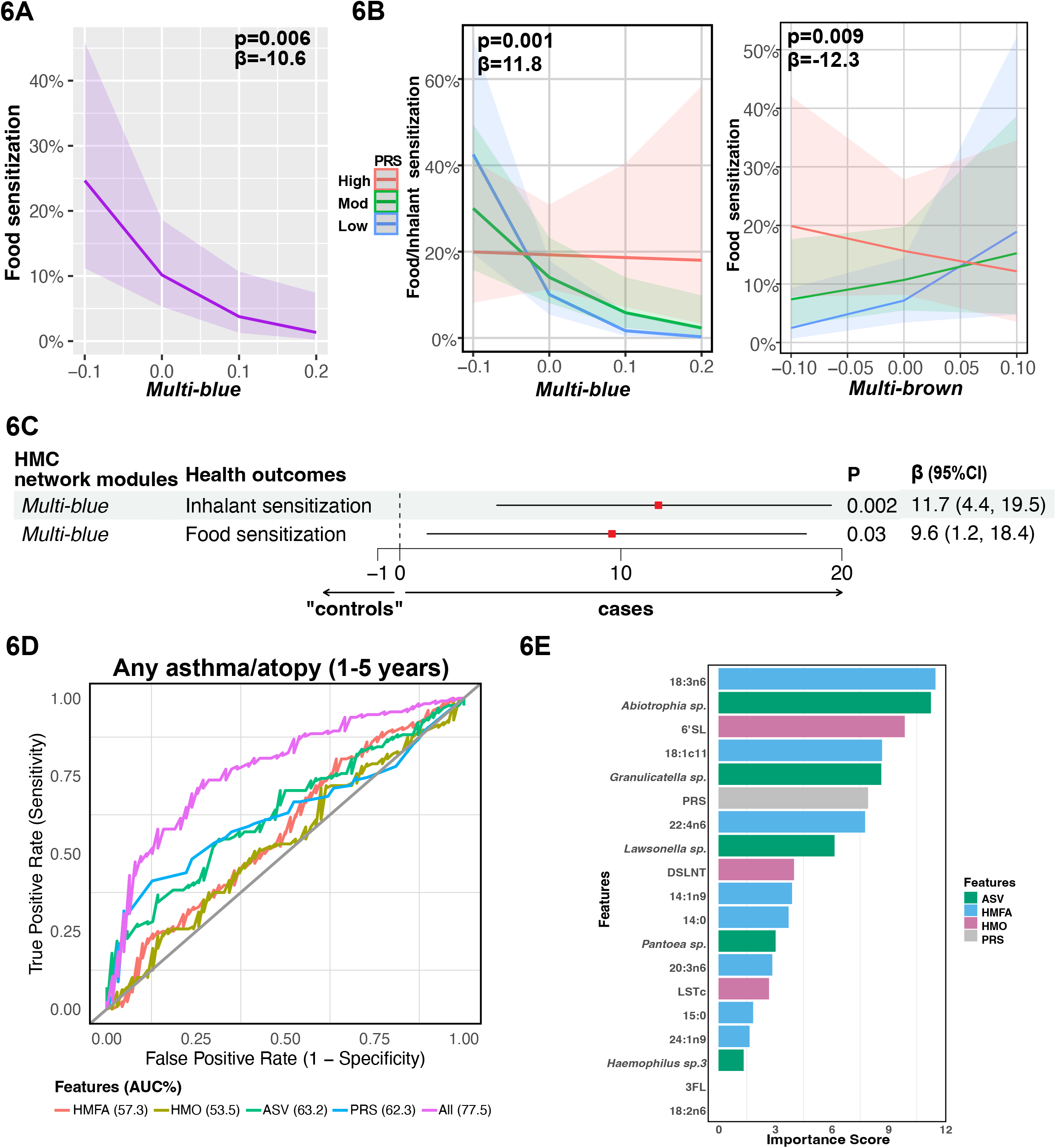
Integration of infant PRS and maternal milk components predicted childhood atopy. Increased abundance of HMCs in the *multi-blue* network module was associated with reduced prevalence of childhood atopy (**A**), among infants with low PRS (**B**, **left**; **C**); increased abundance of HMCs in the *multi-brown* network module was associated with reduced prevalence of childhood atopy among infants with high PRS (**B**, **right**). The interaction beta (β) coefficients in (**B** and **C**) represent how the association between exposure to HMCs and childhood atopy changes with increasing PRS. PRS were categorized into low (< -1 SD), moderate (Mod; -1 to +1 SD), and high (> +1 SD). The GBM model integrating ASVs, HMFAs, HMOs, and infant PRS outperformed models using individual maternal HMC types and infant PRS alone in classifying childhood atopy and related conditions from controls (**D**). Features that best differentiated between cases and “controls” were ranked based on importance scores derived from their contribution to decision tree splits in the GBM (**E**).

### Integration of infant PRS and maternal HMCs improved predictions of childhood atopy

We implemented GBMs to evaluate the individual and combined contributions of all HMCs (i.e., HMFAs, HMOs, and ASVs) with infant PRS in classifying cases vs. “controls”. Given the common co-occurrence of childhood atopy and asthma, we combined multiple case groups, including children with any atopy at ages 1, 3, or 5 years, asthma at age 5 years, or recurrent wheeze between ages 2 and 5 years (n=385). The control group (n=214) consisted of children without any of these conditions (**Table E1**). Feature selection using binomial GLMs identified five ASVs, nine HMFAs, four HMOs, and infant PRS, all of which were significantly associated with childhood atopy in either the main or PRS interaction analyses (**Table E5**). The resulting GBMs integrating all selected features achieved the highest classification performance (AUC=77.5%) in differentiating children with atopy and related conditions from controls (**Figure 6D**). In contrast, GBMs trained and validated on any individual HMC or PRS alone resulted in lower classification performance, with AUCs ranging from 53.5 to 63.2% for the same outcome (**Figure 6D**). We also ranked features (e.g., gamma-linolenic acid (18:3n6), 6′SL, *Abiotrophia sp.*, *Granulicatella sp.*) that best separated cases from “controls” based on importance scores, which quantified their contributions to decision tree splits that improved GBM performance (**Figure 6E)**. Therefore, these results demonstrate that integrating infant genomic susceptibility with multiple types of HMCs improves prediction of childhood atopy compared with models based on individual components.

## Discussion

This study identified interactions between infant polygenic risk and exposure to HMCs associated with childhood atopy risk. We further showed that HMM traits implicated in atopy may be influenced by HMFAs and HMOs, consistent with earlier studies,^5,6,9^ and that co-occurring networks of HMCs were associated with childhood atopy prevalence depending on the infants’ polygenic risk. Importantly, integrating multiple HMC types with infant genomics improved prediction of childhood atopy compared with models based on individual components. Overall, our nuanced study of the mother-milk-infant triad suggests that the composition of human milk modulates disease outcomes among milk-fed infants depending on their individual genomic profiles.

### Interactions between infant PRS and HMM influenced childhood atopy risk

Our PRS×HMM analyses may, in part, explain some of the inconsistent associations previously reported between microbial traits and atopy. For example, we observed that L-histidine biosynthesis (HISTSYN-PWY) in HM was associated with reduced sensitization to inhalant allergens, but only among HM-fed infants with high PRS, and showed no main effect on childhood atopy when PRS was not considered. This protective association, observed only among infants with high PRS, may explain inconsistent findings in earlier studies, which may have included populations with variable genetic risk profiles.^43–45^ Similarly, although *Haemophilus* in the gut and nasopharynx has been linked to both decreased and increased prevalence of allergic sensitization,^46–48^ we found that increased abundance of *Haemophilus* in HM was associated with increased prevalence of food or inhalant sensitization, specifically among infants with high PRS.

We also confirm previously reported associations of Shannon diversity and the HMM network clusters^3^ in HM with childhood atopy. In our PRS×HMM interaction analyses, these associations appeared protective among infants with low or moderate PRS but not among those with high PRS, suggesting that infant genetic susceptibility may modify the effects of microbial diversity. Additionally, *Abiotrophia* was associated with childhood atopy only when infant genomics were considered. We showed that *Abiotrophia* may not confer protective effects among infants with high PRS compared to those with low or moderate PRS. It is possible that *Abiotrophia* fail to counteract, or may even exacerbate, atopic outcomes among children with high PRS. While we observed significant interaction effects with specific HMM traits, we did not identify significant interactions between breastfeeding duration (6 months) and infant genetics. This supports the notion that consideration of breastfeeding as a single homogenous exposure can mask the health benefits of specific milk components (e.g., HMM). Overall, our findings highlight the importance of considering both infant genomics and variations in HMM composition when evaluating early-life risk of atopy.

### Interactions between infant PRS and network clusters of HMCs were associated with childhood atopy

Our multi-component network analysis identified clusters of co-occurring HMM, HMOs, and HMFAs, highlighting that HM functions as an integrated biological system in which components interact rather than act independently. These clusters suggest that HM microbial colonization may be promoted by specific combinations of HMOs and/or HMFAs,^6,49,50^ and likely reflect shared ecological and metabolic relationships that collectively contribute to childhood health benefits. For example, the *multi-brown* cluster was associated with reduced childhood atopy prevalence among HM-fed infants with high genetic risk. This cluster included HMOs such as 2′FL, 6′SL, and LNH, which are known substrates for *Bifidobacterium*.^51^ Other taxa in the same cluster, such as *Lactobacillus*, are indirectly influenced through HMOs-supported cross-feeding interactions with *Bifidobacterium* or through exposure to microbial metabolites.^52,53^ We also identified another *multi-blue* cluster, consisting of *Haemophilus* and *Abiotrophia*, which was associated with reduced childhood atopy prevalence, except among infants with high genetic risk. Notably*, Haemophilus*, *Streptococcus*, and *Veillonella* in the *multi-blue* cluster have been consistently identified in both mothers’ milk and infant gut,^54,55^ supporting the hypothesis^56^ that transfer of these bacteria from HM to the infant gut may influence children’s health outcomes.

### Integration of maternal HMCs and infant PRS improved prediction of childhood atopy

We demonstrate that integrating HMM, HMFAs, HMOs, and infant PRS improves prediction of childhood atopy and related outcomes compared with models based on individual components. The GBM achieved the highest classification performance, supporting the integration of milk-omics exposures with genetic susceptibility. Key contributors to model performance included HMM taxa (e.g., *Granulicatella* and *Abiotrophia*), HMOs (e.g., 6′SL) and HMFAs (e.g., γ-linolenic acid (18:3n6)). Reduced abundance of *Granulicatella* has been observed in asthmatic subjects,^3,57^ and 6′SL supplementation has been shown to reduce food allergy symptoms,^58^ whereas findings on the association between dietary 18:3n6 and AD remains inconsistent.^59^ Notably, 6′SL and 18:3n6 were members of the *multi-brown* network module associated with reduced prevalence of food sensitization among infants with high genetic risk. In summary, the machine learning model trained on all HMCs with PRS differentiated children with atopy and/or asthma from those without these conditions, underscoring the importance of modeling early-life exposures and host genomics in an integrated framework.

### HMOs and HMFAs were associated with atopy-implicated HMM

We observed associations of HMOs and HMFAs with HMM, which contributes to growing evidence of interactions among these bioactive components in HM. It is well known that HMOs are a carbon source that selectively promotes the microbial growth in HM.^5,9,53,60^ Most studies to date have reported the associations of HMOs with *Bifidobacterium*, *Staphylococcus*, and *Lactobacillus*, but the same studies did not investigate the impact of these bacteria on child health.^5,9,53,60^ In this study, we uniquely focused on HMM traits linked to childhood atopy, and our results showed that decreased concentrations of sialylated HMOs such as 6′SL, LSTc, and DSLNH, were associated with increased *Abiotrophia* abundance. Given that *Abiotrophi*a expresses sialidases,^61,62^ the bacteria may hydrolyze α2,6-linked sialic acid from 6′SL and LSTc. It is possible that specific milk taxa selectively use host-derived glycans, thereby shaping HMM structure and subsequent atopy risk.

Compared to HMOs, associations between fatty acids and HMM are less well understood. We observed that HMM traits were associated with the n-6/n-3 PUFA ratio, consistent with findings from an earlier study on rat breast milk.^63^ In our human study, increased abundance of *Abiotrophia* was associated with decreased concentrations of individual n-3 and n-6 PUFAs but increased total n-6/n-3 PUFA and ARA/DHA ratios. This suggests that PUFA ratios, rather than individual PUFAs, play a more important role in these atopy-implicated HMM traits. Earlier studies have reported that prenatal exposure to an increased n-6/n-3 PUFA ratio in maternal blood was associated with reduced prevalence of childhood eczema at ages 6-7 years.^64^ Similarly, in a previous CHILD Study, Miliku *et al.* found that increased ARA/DHA ratio in HM was associated with reduced prevalence of childhood AD and food sensitization among female infants at age 1 year.^11^ Taken together with existing evidence,^11,63,64^ our findings suggest that the nuances of the n-6/n-3 PUFA and ARA/DHA ratios associated with HMM traits may have greater health implications than individual PUFAs alone.

### Limitations and future directions

Although our study leverages deeply phenotyped, longitudinal data from a large sample of mother-infant dyads within the CHILD cohort,^13^ there are several limitations. HM samples were ascertained at a single time point, which limits assessment of temporal variation. Moreover, gene-milk interaction analyses using PRSs cannot identify the specific genes involved, and genome-wide interaction studies could be conducted to further explore interaction effects of individual genetic variants and HMM composition on childhood atopy. This is, however, beyond the scope of our current study. Our machine learning algorithm using GBMs achieved the highest performance in predicting children with atopy and/or asthma-related conditions. This may indicate limited power to classify individual atopy phenotypes due to the limited sample size. Moreover, we acknowledge that our *in silico* study cannot establish causal relationships. Replication in cohorts, with larger sample sizes and experimental studies, is warranted to confirm associations identified in this study and to establish their biological and clinical relevance to childhood atopy. Finally, the use of 16S rRNA sequencing limits taxonomic resolution, and future metagenomic and/or culture-based studies are needed to investigate HMM at the species and strain levels.

## Conclusion

This study identifies gene-milk interactions linking infant genomics and exposure to HMCs to childhood atopy. Integration of maternal HMOs, HMFAs, and HMM with infant PRS improves prediction of childhood atopy. These findings support the potential for personalized interventions, such as specific probiotics or synbiotics, to help prevent childhood atopy. For example, probiotics enriched with taxa from the *multi-brown* (e.g., *Bifidobacterium longum*) and *multi-blue* (e.g., *Abiotrophia*) network clusters could be prioritized for infants with high and low PRS, respectively. Synbiotic formulations may also incorporate HMOs such as 6′SL in the *multi-brown* cluster for infants with high PRS and 3FL in the *multi-blue* cluster for those with low PRS.

## Acknowledgments

We thank Kelsey Fehr and Shirin Moossavi for their work in processing and analyzing the HMM data. We are grateful to all the CHILD families who took part in this study, and the whole CHILD team, which includes interviewers, nurses, computer and laboratory technicians, clerical workers, research scientists, volunteers, managers, and receptionists. For a list of investigators and enrolling centers visit www.childcohort.ca. Computational analyses were performed on resources and with support provided by the Centre for Advanced Computing (CAC) at Queen’s University in Kingston, Ontario.

## Data Availability

The demultiplexed 16S rRNA gene sequencing data have been previously deposited into the Sequence Read Archive (SRA) under the BioProject accession numbers (NCBI) PRJNA481046 and PRJNA597997 with SRA number SRP153543.

## Abbreviations

AD: Atopic dermatitis
ARA: Arachidonic acid
ASV: Amplicon sequence variant
AUC: Area under the curve
CLR: Centered-log-ratio
DHA: Docosahexaenoic acid
FA: Fatty acid
GBM: Gradient-boosting machine
GLM: Generalized linear model
HM: Human milk
HMC: Human milk component
HMFA: Human milk fatty acid
HMM: Human milk microbiota
HMO: Human milk oligosaccharide
n-6: omega-6
n-3: omega-3
PC: Principal component
PRS: Polygenic risk score
PUFA: Polyunsaturated fatty acid
QC: Quality control
SD: Standard deviation
SPT: Skin prick test
WGCNA: Weighted gene correlation network analysis

