## Supplementary Information for "Interactions between human milk components and infant polygenic risk predict childhood atopy"

### Supplementary Methods

#### Childhood atopy and related outcomes

Childhood atopy is defined as the presence of atopic dermatitis (AD)[1,2] and/or sensitization to food or inhalant allergens (i.e., positive skin prick tests (SPTs) at ages 1, 3, and 5 years (**Table E1**).[3] As previously described, children were diagnosed with AD during the scheduled visits, based on a careful assessment of clinical history and examinations, aligning with the criteria derived from the UK working party document.[1,2] A SPT response to 4 common food or 13 inhalant allergens (peanut, milk, egg white, and soy and Alternaria tenuis, cat hair, dog epithelium, house dust mite [Dermatophagoides pteronyssinus and Dermatophagoides farinae], cockroach, Penicillium, Cladosporium, Aspergillus fumigatus, tree, grass, weed, and ragweed) was defined as a wheal diameter of ≥ 2mm.[3] Glycerin and histamine were used as the negative and positive controls, respectively.[3]

The secondary outcome included children with atopy- or asthma-related conditions (**Table E1**). The case group included 64 children diagnosed with definite asthma at age 5 years and 140 children who experienced recurrent wheeze between ages 2 and 5 years. Specifically, definite asthma was diagnosed by an experienced allergist or asthma specialist at age 5.[4] As previously described, wheezing symptoms were assessed by expert physicians at ages 3 and 5 years and standardized questionnaires between ages 2 and 5 years. Specifically, parents completed repeated questionnaires on their children's wheeze frequency and triggers at 5 time points (24, 30, 36, 48, and 60 months postpartum).

#### Human milk microbiota (HMM) analyses

The two batches of HMM samples were collected in 2016 and 2019, with the second batch including a subset of lactating mothers enriched for maternal and infant chronic conditions (i.e., obesity, asthma, and atopy).[5,6] HMM was analyzed from 885 mothers (**Figure E1**), with genomic

DNA extracted from 1 mL of their milk samples using the Zymo Research Quick-DNA Fungal/Bacterial DNA kit, as previously described.[7,8] Within these samples, the V4 hypervariable region of the 16S rRNA gene was amplified and sequenced using modified F515/R806 primers on a MiSeq platform (Illumina, San Diego, CA, USA).[9,10] To prepare the sequencing library, sterile DNA-free water was used as a negative control, and a mock community was previously created by extracting DNA from eight species with known theoretical relative abundances (Zymo Research, USA) and was included as the positive control. Data preprocessing of HMM has been previously described.[8,11] Briefly, overlapping paired-end reads were merged and processed following the standard DADA2 denoising pipeline from QIIME 2 v.2019.10 (https://qiime2.org).[9,12,13] Qualified and trimmed reads were taxonomically annotated with SILVA v138 reference database at 99% sequence similarity. To identify and remove potential sequencing contaminants and artifacts,[14,15] a two-stage process was employed using the Decontam[16] and Phyloseq[17] R packages. Additional batch-specific contaminants were removed based on the inherent structure of the HMM dataset.[18] Spurious ASVs with <0.001% reads per sample on average and samples with <8,000 sequencing reads were eliminated.[19] This resulted in 2,641 ASVs remaining across 885 samples.

To infer potential common metabolic pathways contributed by milk ASVs, we previously applied PICRUSt2[20] to predict the MetaCyc[21] metabolic pathway abundances based on 16S rRNA gene sequences.[11] Abundances of ASVs and metabolic pathways were centered-log-ratio (CLR) transformed using the CoDaSeq R package,[22] after replacing zeros with imputed values by applying the Geometric Bayesian multiplicative method in the zComposition R package.[11,23] Following the rarefaction of read count data to a minimum depth of 8,005 reads, alpha-diversity (i.e., Shannon) was computed using the Phyloseq R package.[11,17] Previously, we conducted a weighted gene co-expression network analysis (WGCNA)[24] using a 'signed' network model, which clustered 179 CLR-transformed ASVs into six co-abundance network modules with potentially shared biological processes.[11]

**Human milk fatty acid (HMFA) and oligosaccharide (HMO) analyses**

The following 28 HMFAs were previously identified based on commercial standards and reported as relative proportions of total fatty acids: capric acid (10:0), lauric acid (12:0), myristic acid (14:0), physeteric acid (14:1n9), pentadecylic acid (15:0), palmitic acid (16:0), 7-hexadecenoic acid (16:1n9), heptadecanoic acid (17:0), stearic acid (18:0), trans-vaccenic acid (TVA; 18:1t11), oleic acid (18:1n9), cis-vaccenic acid (18:1c11), linoleic acid (LA; 18:2n6), arachidic acid (20:0), γ-linolenic acid (GLA; 18:3n6), α-linolenic acid (ALA; 18:3n3), eicosadienoic acid (20:2n6), dihomo-γ-linolenic acid (DGLA; 20:3n6), arachidonic acid (ARA; 20:4n6), conjugated linoleic acid (CLA; a family of linoleic acid isomers), eicosapentaenoic acid (EPA; 20:5n3), nervonic acid (24:1n9), docosahexaenoic acid (DHA; 22:6n3), lignoceric acid (24:0), osbond acid (22:5n6), docosapentaenoic acid (DPA; 22:5n3), eicosatetraenoic acid (20:4n3), and adrenic acid (22:4n6) (**Table E2**).[25]

HMOs were quantified in nmol/mL of human milk by comparing their retention times with commercial standards and using mass spectrometry analysis (**Table E3**).[25] The 19 HMOs analyzed included: 2′-fucosyllactose (2′FL), 3-fucosyllactose (3FL), 3′-sialyllactose (3′SL), 6′-sialyllactose (6′SL), lacto-N-tetraose (LNT), lacto-N-neotetraose (LNnT), lacto-N-fucopentaose 1-3 (LNFP1-3), sialyl-LNTb (LSTb), sialyl-LNTc (LSTc), difucosyl-LNT (DFLNT), disialyl-LNT (DSLNT), fucosyl-lacto-N-hexaose (FLNH), difucosyl-lacto-N-hexaose (DFLNH), fucosyl-disialyl-lacto-N-hexaose (FDSLNH), disialyl-lacto-N-hexaose (DSLNH), difucosyllactose (DFLac), and lacto-N-hexaose (LNH) (**Table E3**).[25]

**Infant genomics**

A total of 557,005 single nucleotide variants (SNVs) were genotyped on the Illumina HumanCoreExome BeadChip after isolating DNA from the cord blood of 2,967 infants (**Figure E1**). Quality control (QC) steps were performed previously using the PLINK software.[26–28] Briefly, subjects with sex discrepancies were removed, determined as females with X chromosome F-statistics >0.2 and males with Y chromosome F-statistics <0.8. Furthermore, subjects subjects with values exceeding ±2 standard deviations of non-homozygous variants, >10% genotype missingness, or >0.185 inbreeding coefficient (i.e., to assess relatedness) were omitted. Following the removal of SNVs with missingness >5%, the remaining 551,033 SNVs were imputed in 2,835 infants using the Haplotype Reference Consortium (HRC; r1.1 2016) data on the Michigan server.[29]

**Polygenic risk score (PRS) Calculation and Evaluation**

To capture the cumulative effects of genetic variants on atopy, we computed PRS for atopic outcomes in the CHILD dataset using the Polygenic Score Catalog Calculator (pgsc_calc v2.0.0).[30] PRS calculation is a widely used because the combined effect of genetic variants typically provides greater statistical power than an individual locus for predicting disease outcomes with polygenic origins. Prior to calculating PRS, we removed strand-ambiguous, multiallelic, and duplicated variants using the default settings of the *pgsc_calc* tool.[30] Following these QC steps, we imputed PRS weights for atopic outcomes in the CHILD dataset[4] by leveraging existing PRS weights from the PGS Catalog (https://www.pgscatalog.org/), which represent the magnitude of association between genetic variants and atopy. Specifically, we used PRS weights reported by Tanigawa *et al.*[31] (PGS ID: PGS001285) from 7,312 single nucleotide polymorphisms (SNPs) in 245,788 subjects with physician-diagnosed allergic diseases (i.e., hay fever, rhinitis, and/or eczema).

To evaluate the predictive performance of our constructed PRS for atopic outcomes in CHILD,[4] we assessed its association with childhood atopy between ages 1 and 5 years in a binomial generalized linear model (GLM), adjusted for study center, infant sex, antibiotic exposure during the first year of life, and ethnicity reported by parents using questionnaires. Other standard metrics, including Nagelkerke's pseudo $R^2$ and area under the curve (AUC), were applied to further evaluate the predictive performance of our constructed PRS for atopic outcomes between ages 1-5 in CHILD.[4]

### Clustering of correlated HMOs and HMFAs

Given that individual HMOs and HMFAs may be correlated with each other, Bonferroni correction was applied based on the number of HMO and HMFA clusters estimated. The six clusters of correlated HMOs were previously determined using Pearson correlation and hierarchical clustering of the 19 individual HMOs.[26] We applied the same method to group the 28 individual HMFAs into clusters of correlated HMFAs.[26] (**Figure E2**) The numbers of clusters were optimally determined using the silhouette method in the factoextra[32] and dendextend[33] R packages.

### Multi-component network analysis

We conducted multi-component network analyses of 179 CLR-transformed ASVs, 19 19 rank-based inverse normal–transformed HMOs, and 28 rank-based inverse normal–transformed HMFAs using the WGCNA R package.[24] Prior to the network analysis, all HM features were scaled to a common range of –2 to +2 to ensure equal weight for HMM, HMOs, and HMFAs in the correlation calculations, while preserving the same distribution of values within each subject. Following the framework of a 'signed' WGCNA,[24] an adjacency matrix was constructed based on the correlation between each pair of HM features. The optimal soft thresholding power (β) was chosen at 7 with a scale-free topology fitting index ($R^2$) reaching 0.85 (**Figure E3A**). Hierarchical clustering and dynamic tree cut methods, with a minimum cluster size of 10, were employed to generate modules of correlated HMCs (**Figure E3B**). These modules, labelled by colors, were visualized using the Cytoscape software[34] and subsequently subjected to principal component (PC) analysis to compute an eigenvalue for each module. Each module eigenvalue, defined as the first PC, was used to optimally summarize the co-occurrence patterns of all HMCs within the module.[24]

### Feature selection in machine learning

Prior to training the gradient-boosting machine (GBM), all features were standardized to a mean of 0 and a standard deviation of 1 to equalize variance and prevent features with larger natural spreads from disproportionately influencing model performance. We then performed feature selection by testing both the main effects of exposure to HMOs and HMFAs and their interactions with PRS on childhood atopy using binomial GLM-based regression models (**Table E5**). These models adjusted for the same covariates used in the PRS×HMM analyses: study center, milk batch, time from milk sample collection to processing, infant sex, antibiotic exposure during the first year of life, and infant ethnicity reported by parents using questionnaires.

### Supplementary Results

#### Infant PRS was associated with childhood atopic outcomes

The PRS constructed using the PGS Catalog (https://www.pgscatalog.org/) was significantly associated with all atopic-related outcomes at ages 1-5 years in the CHILD Cohort Study (**Figure E4A-H**). Specifically, infant PRS was also associated with childhood atopic dermatitis (AD; P=1.1E-6, β=0.5) (**Figure E4A and E**), positive SPTs (P=2.5E-5, β=0.4) (**Figure E4B and F**), inhalant sensitization (P=1.1E-4, β=0.4) (**Figure E4C and G**), and food sensitization (P=1.5E-4, β=0.4) (**Figure E4D and H**) in the CHILD Cohort Study.

#### Identification of clusters of correlated HMFAs

We applied Pearson correlation and hierarchical clustering to group the 28 individual HMFAs into three clusters of correlated HMFAs (**Figure E2**).[26] To account for multiple tests in GLM analyses assessing associations between the 28 HMFAs and HMM, Bonferroni correction was applied based on the three HMFA clusters estimated (**Figure E2**).

#### Identification of co-occurring network clusters of HMFAs, HMOs, and HMM.

Previously, we conducted WGCNA[24] using a 'signed' network model, which clustered 179 CLR-transformed ASVs into six co-abundance network modules with potentially shared biological processes.[11] In this manuscript, to study the HM as a biological system, multi-component network analyses of 179 ASVs, 19 HMOs, and 28 HMFAs using the WGCNA R package[24] were clustered into seven network modules (**Table E8**). Four of the seven modules consisted of members from only one HMC type (**Table E8**). The *multi-red*, *multi-blue*, and *multi-brown* modules, however, were made up of more than one HMC type (**Table E8**). For example, the *multi-blue* module consisted of 4 HMOs and 45 ASVs, whereas the *multi-brown* module consisted of 19 HMFAs, 9 HMOs, and 11 ASVs.

**Features selected for machine learning analyses**

In addition to determining that exposure to five human milk taxa were associated with childhood atopic outcomes, our feature selection steps also identified four HMOs and nine HMFAs that were associated with childhood atopy, with or without considering infant PRS after Bonferroni multiple testing correction (**Table E5**). In our main association analyses without considering infant PRS, decreased concentrations of two HMOs (DSLNT and 3FL) and three HMFAs (i.e., 14:0, 15:0, and 14:1n9) were associated with reduced prevalence of childhood sensitization to inhalant or food allergens between ages 1 and 5 years ($P_{Bonf}$=6.8E-03, β=-0.4). Increased concentrations of four HMFAs were also associated with increased prevalence of allergic sensitization to inhalant or food allergens between ages 1 and 5 years (P=6.8E-03, β=0.8). In our interaction analyses between infant PRS and HMOs, increased concentrations of 18:3n6 was associated with increased prevalence of allergic sensitization to food or inhalant allergens at age 1 year among HM-fed infants with high PRS, and the same exposure was associated with reduced prevalence of childhood allergic sensitization at age 1 year among infants with low PRS ($P_{Bonf}$=0.03, β=0.3). Moreover, increased concentrations of 24:1n9, LSTc, and 6'SL were associated with reduced allergic sensitization to food or inhalant allergens between ages 1 and 5 years among infants with high PRS ($P_{Bonf}$=0.02, β=-0.5). Among infants with low PRS, exposure to increased concentrations of 24:1n9, LSTc, and 6'SL were associated with increased allergic sensitization to food or inhalant allergens between ages 1 and 5 years ($P_{Bonf}$=0.02, β=-0.5).

### List of Supplementary Tables and Figures

**Table E1:** Characteristics of 689 mother-infant dyads (sub-cohort) included in this study compared with all dyads (n=3,542) from the full CHILD Cohort Study.

**Table E2.** Summary of the human milk fatty acids (HMFAs) quantified in 1,200 mothers of the CHILD Cohort Study.

**Table E3.** Summary of the human milk oligosaccharides (HMOs) quantified in 1,206 mothers of the CHILD Cohort Study.

**Table E4:** Main effects of human milk microbiota (HMM) traits[11] and their interactions with infant PRS were associated with childhood atopy at ages 1-5 years. These atopy-implicated HMM traits were carried forward for downstream association analyses with HMOs and HMFAs.

**Table E5:** Main effects of maternal HMM,[11] HMFAs, and HMOs and their interactions with infant PRS, were associated with childhood atopy. These maternal milk components and infant PRS were selected to train gradient boosting tree models to classify childhood atopy/asthma versus controls.

**Table E6:** HMFAs were associated with atopy-implicated HMM traits.

**Table E7:** HMOs were associated with atopy-implicated HMM traits.

**Table E8:** Weighted gene co-expression network analysis (WGCNA) clustered 179 individual milk ASVs, 19 individual HMOs, and 28 HMFAs into seven multi-component network modules encompassing 226 milk features: 10 in multi-black, 49 in multi-blue, 39 in multi-brown, 32 in multi-purple, 15 in multi-red, 32 in multi-orange, and 49 in multi-turquoise.

**Figure E1:** Study flowchart of maternal milk sample and infant genomic analyses.

**Figure E2:** Clustering of 28 individual HMFA profiles among 1,200 lactating mothers, related to Methods.

**Figure E3:** WGCNA clustered 179 amplicon sequence variants (ASVs), 19 human milk oligosaccharides (HMOs), and 28 human milk fatty acids (HMFAs) into seven co-occurring network modules.

**Figure E4:** Polygenic risk scores (PRS) were associated with childhood atopy at ages 1-5 years in the CHILD dataset.

**Supplementary Tables**

**Table E1:** Characteristics of 689 mother-infant dyads (sub-cohort) included in this study compared with all dyads (n=3,542) from the full CHILD Cohort Study.

| | Variables | Mean ± SD or n (%)[a] | |
|---|---|---|---|
| | | Mother-infant dyads (n = 689) | All eligible dyads (n = 3,542) |
| Maternal | **Age (years)** | 33 ± 4.2 | 32.3 ± 4.7 |
| | **Reported ethnicity** | | |
| | Central European (CEU) | 518 (75.2) | 2359 (72.9) |
| | Other | 171 (24.8) | 876 (27.1) |
| | **Milk microbiome batches** | | |
| | Batch 1 | 292 (42.4) | 344 (38.9) |
| | Batch 2 | 397 (57.6) | 541 (61.1) |
| | **Time (in seconds) between sample collection at home and processing** | 68229 ± 70844 | 108835 ± 456953 |
| | **Annual total household income (above $100,000 Canadian dollars)** | 351 (56.7) | 1504 (52.6) |
| | **Have a post-secondary degree** | 556 (81.3) | 2407 (76.3) |
| | **Breastfeeding for ≥6 months** | 689 (100) | 2322 (76.2) |
| | **Study centers** | | |
| | Edmonton | 142 (20.6) | 780 (23.7) |
| | Toronto | 195 (28.3) | 778 (23.6) |
| | Edmonton | 179 (26) | 742 (22.5) |
| | Toronto | 173 (25.1) | 996 (30.2) |
| Children | **Antibiotics use at age 1 year** | 135 (19.6) | 600 (18.4) |
| | **Reported ethnicity** | | |
| | Central European (CEU) | 450 (65.3) | 2061 (64) |
| | Others | 239 (34.7) | 1159 (36) |
| | **Sex** | | |
| | Female | 310 (45) | 1642 (47.4) |
| | Male | 379 (55) | 1820 (52.6) |
| | **Atopy** | | |
| | Any atopic dermatitis | | |
| | at 1 year | 105 (15.5) | 349 (25.2) |
| | at 3 years | 100 (16.3) | 327 (24) |
| | at 5 years | 117 (19.3) | 367 (26.2) |
| | between 1 to 5 years | 206 (34.6) | 707 (40.6) |
| | between 3 to 5 years | 164 (28.2) | 530 (33.9) |
| | Any food/inhalant sensitization (i.e., positive skin prick tests (SPTs)) | | |
| | at 1 years | 117 (17.5) | 418 (28.8) |
| | at 3 years | 122 (18.9) | 408 (28.3) |
| | at 5 years | 137 (22/5) | 524 (33.6) |
| | at 1 to 5 years | 239 (39.3) | 888 (46.2) |
| | at 3 to 5 years | 188 (31.2) | 695 (40.2) |
| | Any food sensitization | | |
| | at 1 years | 99 (14.9) | 334 (24.4) |

| | | | |
|---|---|---|---|
| | at 3 years | 50 (7.8) | 167 (13.9) |
| | at 5 years | 40 (6.6) | 143 (12.1) |
| | at 1 to 5 years | 120 (20.3) | 414 (28.6) |
| | at 3 to 5 years | 64 (10.9) | 214 (17.1) |
| | Any inhalant sensitization | | |
| | at 1 years | 26 (4.9) | 124 (10.7) |
| | at 3 years | 95 (14.7) | 327 (24) |
| | at 5 years | 125 (20.5) | 469 (31.2) |
| | at 1 to 5 years | 179 (29.9) | 691 (40) |
| | at 3 to 5 years | 165 (27.5) | 622 (37.6) |
| | **Definite asthma at age 5 years** | 64 (11.1) | 171 (6.8) |
| | **Recurrent wheeze at ages 2 to 5 years** | 140 (20.8) | 444 (17.5) |
| | **Any atopy at ages 1 to 5 years** | 333 (55.7) | 1253 (50.6) |
| | **Any atopy, asthma, and/or recurrent wheeze at ages 1 to 5 years**[b] | 385 (64.3) | 1458 (58.5) |

[a]Percentages reflect the proportion of non-missing data.
[b]Case groups were compared with 214 controls among 689 dyads (sub-cohort) or 1,034 controls among 3,542 dyads from the full cohort. Controls consisted of children without any asthma, recurrent wheeze, atopy (including food or inhalant allergies), or atopic dermatitis (AD) at ages 1-5 years.

**Table E2.** Summary of the human milk fatty acids (HMFAs) quantified in 1,200 mothers of the CHILD Cohort Study.

| HMFA Abbreviation | HMFA Full Name | Mean | Standard Deviation | Minimum | Maximum |
|---|---|---|---|---|---|
| **Total SFA %** | | | | | |
| 10:0 | Capric acid | 0.7 | 0.3 | 0.006 | 2 |
| 12:0 | Lauric acid | 4.8 | 1.7 | 1.2 | 15.8 |
| 14:0 | Myristic acid | 6 | 1.9 | 2.1 | 18.5 |
| 16:0 | Palmitic acid | 20.9 | 2.9 | 13.2 | 36.1 |
| 17:0 | Heptadecanoic acid | 0.3 | 0.08 | 0.1 | 0.6 |
| 18:0 | Stearic acid | 6.5 | 1.4 | 2.6 | 18.7 |
| 20:0 | Arachidic acid | 0.2 | 0.08 | 0.02 | 2 |
| 24:0 | Lignoceric acid | 0.05 | 0.03 | 0.002 | 0.3 |
| **Total MUFA %** | | | | | |
| 14:1n9 | Physeteric acid | 0.2 | 0.09 | 0.02 | 0.7 |
| 16:1n9 | 7-Hexadecenoic acid | 2.7 | 0.7 | 0.8 | 5.2 |
| 18:1n9 | Oleic acid | 37 | 3.8 | 20.3 | 52.5 |
| CLA | Conjugated linoleic acid | 0.02 | 0.01 | 0.001 | 0.2 |
| 24:1n9 | Nervonic acid | 0.05 | 0.02 | 0.002 | 0.2 |
| TVA | Trans-vaccenic acid | 1.5 | 1.2 | 0.1 | 7.4 |
| 18:1C11 | Cis-vaccenic acid | 1.7 | 0.5 | 0.1 | 6.2 |
| **Total PUFA%** | | | | | |
| 18:2n6 | Linoleic acid (LA) | 13.6 | 3.1 | 6.2 | 27 |
| 18:3n6 | γ-Linolenic acid (GLA) | 0.1 | 0.06 | 0.02 | 0.5 |
| 20:2n6 | Eicosadienoic acid | 0.2 | 0.06 | 0.07 | 1.3 |
| 20:3n6 | Dihomo-γ-linolenic acid (DGLA) | 0.3 | 0.1 | 0.06 | 0.8 |
| 20:4n6 | Arachidonic acid (ARA) | 0.4 | 0.1 | 0.08 | 0.8 |
| 22:4n6 | Adrenic acid | 0.04 | 0.03 | 0.001 | 0.1 |
| 22:5n6 | Osbond acid | 0.03 | 0.01 | 0.005 | 0.1 |
| 18:3n3 | α-Linolenic acid (ALA) | 1.9 | 0.7 | 0.08 | 5.8 |
| 20:4n3 | Eicosatetraenoic acid | 0.08 | 0.03 | 0.01 | 0.4 |
| 20:5n3 | Eicosapentaenoic acid (EPA) | 0.09 | 0.08 | 0.01 | 1.1 |
| 22:5n3 | Docosapentaenoic acid (DPA) | 0.1 | 0.06 | 0.04 | 0.5 |
| 22:6n3 | Docosahexaenoic acid (DHA) | 0.2 | 0.2 | 0.02 | 1.5 |
| **Total HMFAs and ratios** | | | | | |
| Total SFAs | Total saturated fatty acids | 39.8 | 5.2 | 24.5 | 59.9 |
| Total MUFAs | Total mono unsaturated fatty acids | 43 | 3.8 | 25.8 | 57.2 |
| n-6 | Total omega-6 | 14.8 | 3.2 | 7.1 | 28.1 |
| n-3 | Total omega-3 | 2.4 | 0.8 | 0.6 | 7.2 |

| | | | | | |
|---|---|---|---|---|---|
| Total PUFAs | Total polyunsaturated fatty acids | 17.2 | 3.7 | 8.5 | 32.2 |
| ARA/DHA | Arachidonic to DHA ratio | 2.6 | 1.5 | 0.3 | 17.6 |
| Total ARA/DHA and EPA | Arachidonic to (DHA and EPA) ratio | 1.8 | 0.8 | 0.2 | 5.9 |
| Total n-6/n-3 | Omega-6 to omega-3 PUFAs ratio | 6.5 | 1.9 | 1.8 | 23.2 |

**Table E3.** Summary of the human milk oligosaccharides (HMOs) quantified in 1,206 mothers of the CHILD Cohort Study.

| HMO Abbreviation | HMO Full Name | Mean (nmol/ml) | Standard Deviation | Minimum | Maximum |
|---|---|---|---|---|---|
| 2'FL | 2'-fucosyllactose | 4828 | 3606 | 0 | 17514 |
| 3FL | 3-fucosyllactose | 461 | 304 | 9 | 2971 |
| 3'SL | 3'-sialyllactose | 582 | 380 | 20 | 6640 |
| 6'SL | 6'-sialyllactose | 318 | 218 | 34 | 2704 |
| DFLac | Difucosyllactose | 471 | 389 | 2 | 3392 |
| DFLNH | Difucosyllacto-N-hexaose | 51 | 60 | 3 | 585 |
| DFLNT | Difucosyllacto-N-tetrose | 1364 | 743 | 16 | 3682 |
| DSLNH | Disialyllacto-N-hexaose | 88 | 63 | 2 | 567 |
| DSLNT | Disialyllacto-N-tetraose | 264 | 164 | 15 | 1084 |
| FDSLNH | Fucodisialyllacto-N-hexaose | 368 | 236 | 4 | 1705 |
| FLNH | Fucosyllacto-N-hexaose | 59 | 48 | 1 | 375 |
| LNFP1 | Lacto-N-fucopentaose-I | 909 | 889 | 12 | 5588 |
| LNFP2 | Lacto-N-fucopentaose-II | 2022 | 966 | 66 | 5578 |
| LNFP3 | Lacto-N-fucopentaose-III | 88 | 52 | 13 | 332 |
| LNH | Lacto-N-hexaose | 73 | 42 | 0 | 296 |
| LNnT | Lacto-N-neotetraose | 666 | 459 | 0 | 4055 |
| LNT | Lacto-N-tetrose | 1239 | 651 | 64 | 5559 |
| LSTb | Sialyl-lacto-N-tetraose b | 115 | 69 | 9 | 596 |
| LSTc | Sialyl-lacto-N-tetraose c | 70 | 63 | 1 | 817 |
| Sia | Sialic acid bound HMOs | 2526 | 806 | 675 | 8179 |
| Fuc | Fucose bound HMOs | 12507 | 4706 | 1802 | 23291 |
| SUM | Total HMO Concentrations | 14036 | 3955 | 4761 | 25066 |

**Table E4:** Main effects of human milk microbiota (HMM) traits[11] and their interactions with infant PRS were associated with childhood atopy at ages 1-5 years. These atopy-implicated HMM traits were carried forward for downstream association analyses with HMOs and HMFAs.

| HMM traits | HMM or PRS×HMM | Any atopic outcomes at ages 1-5 years | Estimates for HMM | P values[a] | $P_{Bonf}$[b] |
|---|---|---|---|---|---|
| **CLR-transformed ASV abundance** | | | | | |
| *Haemophilus sp. 3* | PRS×HMM | Food/inhalant | 0.2 | 4.6E-05 | 3.9E-03 |
| *Abiotrophia sp* | PRS×HMM | Inhalant | 0.3 | 6.4E-05 | 5.5E-03 |
| *Granulicatella sp.* | HMM | food | -0.2 | 4.7E-04 | 0.04 |
| **CLR-transformed MetaCyc Pathway abundance** | | | | | |
| Superpathway of heme biosynthesis from glutamate (PWY-5918) | PRS×HMM | Inhalant | -0.5 | 2.1E-04 | 0.02 |
| L-ornithine biosynthesis (GLUTORN-PWY) | PRS×HMM | Inhalant | -0.4 | 2.6E-04 | 0.03 |
| preQ0 biosynthesis (PWY-6703) | PRS×HMM | Inhalant | -0.4 | 3.4E-04 | 0.03 |
| L-histidine biosynthesis (HISTSYN-PWY) | PRS×HMM | Inhalant | -0.4 | 4.20E-04 | 0.04 |
| Partial TCA cycle (obligate autotrophs) (PWY-5913) | PRS×HMM | ad | -0.5 | 4.9E-04 | 0.05 |
| Mixed acid fermentation (FERMENTATION-PWY) | PRS×HMM | ad | -0.5 | 5.4E-04 | 0.05 |
| **HMM network clusters** | | | | | |
| *Green* | PRS×HMM | Food/inhalant | 12.8 | 1.20E-03 | N/A |
| *Yellow* | HMM | Food/inhalant | -8.3 | 2.8E-03 | N/A |
| *Yellow* | HMM | ad | -8.5 | 3.3E-03 | N/A |
| *Yellow* | HMM | inhalant | -8.6 | 5.1E-03 | N/A |
| *Green* | PRSxHMM | inhalant | 11.2 | 6.9E-03 | N/A |
| *Yellow* | HMM | food | -8.9 | 0.01 | N/A |
| *Green* | HMM | Food/inhalant | -6.9 | 0.01 | N/A |
| *Green* | HMM | food | -8.9 | 0.02 | N/A |
| *Turquoise* | HMM | ad | 6 | 0.04 | N/A |
| *Green* | HMM | inhalant | -6.3 | 0.05 | N/A |
| *Turquoise* | HMM | Food/inhalant | 5.6 | 0.05 | N/A |
| *Yellow* | PRSxHMM | inhalant | 7.4 | 0.05 | N/A |
| *Green* | PRSxHMM | ad | 7.2 | 0.05 | N/A |
| **Alpha-diversity** | | | | | |
| Shannon | HMM | Food/inhalant | -0.4 | 5.6E-03 | N/A |
| Shannon | PRS×HMM | inhalant | 0.4 | 0.01 | N/A |
| Shannon | HMM | inhalant | -0.4 | 0.01 | N/A |

| | | | | | |
|---|---|---|---|---|---|
| Shannon | HMM | ad | -0.3 | 0.03 | N/A |
| Shannon | HMM | food | -0.4 | 0.03 | N/A |
| Shannon | PRS×HMM | Food/inhalant | 0.4 | 0.03 | N/A |

[a]Unadjusted P values were reported for Shannon diversity and network clusters.
[b]P values for 179 individual amplicon sequence variants (ASVs) and 398 individual metabolic pathways were adjusted using Bonferroni correction for 85 and 98 independent tests, respectively.

**Table E5:** Main effects of maternal HMM,[11] HMFAs, and HMOs and their interactions with infant PRS, were associated with childhood atopy. These maternal milk components and infant PRS were selected to train gradient boosting machines to classify childhood atopy/asthma versus controls.

| HMM traits | HM or PRS×HM | Associated atopic outcomes | Estimate for HMM | Top P value | Top $P_{Bonferroni}$[a] |
|---|---|---|---|---|---|
| **CLR-transformed ASV abundance** | | | | | |
| *Haemophilus sp. 3* | PRS×HM | food/inhalant1to5y | 0.2 | 4.6E-05 | 3.9E-03 |
| *Abiotrophia sp* | PRS×HM | food/inhalant3to5y<br>inhalant1to5y; inhalant3to5y | 0.3 | 6.4E-05 | 5.5E-03 |
| *Lawsonella sp.* | HM | food/inhalant5y;<br>food5y; food3to5y<br>inhalant5y | 0.2 | 3.5E-04 | 0.03 |
| *Granulicatella sp.* | HM | food/inhalant1y;<br>food1to5y | -0.2 | 4.7E-04 | 0.04 |
| *Pantoea sp.* | PRS×HM | inhalant3y | -0.6 | 5.2E-04 | 0.04 |
| **Rank-based inverse normal transformed HMFAs** | | | | | |
| 22:4n6 | HM | food5y | 0.8 | 1.2E-03 | 3.7E-03 |
| 18:2n6 | HM | inhalant1y | 0.6 | 3.6E-03 | 0.01 |
| 18:1c11 | HM | Food/inhalant3to5y; Food/inhalant5y;<br>inhalant3to5y; inhalant5y; | 0.3 | 4.2E-03 | 0.01 |
| 20:3n6 | HM | food5y | 0.6 | 0.01 | 0.03 |
| 14:0 | HM | inhalant1y | -0.5 | 0.01 | 0.03 |
| 18:3n6 | PRSxHM | Food/inhalant1y | 0.3 | 0.01 | 0.03 |
| 15:0 | HM | Food/inhalant3y;<br>inhalant1y; inhalant1to5y | -0.5 | 0.02 | 0.05 |
| 14:1n9 | HM | inhalant1y | -0.5 | 0.02 | 0.05 |
| 24:1n9 | PRSxHM | food3y;food3to5y | -0.4 | 0.02 | 0.05 |
| **Rank-based inverse normal transformed HMOs** | | | | | |
| DSLNT | HM | Food/inhalant5y;<br>inhalant1to5y; inhalant3to5y; inhalant5y | -0.4 | 1.1E-03 | 6.8E-03 |
| 3FL | HM | inhalant1y | -0.7 | 4.7E-03 | 0.03 |

| | | | | | |
|---|---|---|---|---|---|
| LSTc | PRS×HM | food1to5y | -0.4 | 3.1E-03 | 0.02 |
| 6’SL | PRS×HM | food3y | -0.5 | 6.8E-03 | 0.04 |

[a]HM traits associated with childhood asthma and atopy ($P_{Bonferroni} < 0.05$).

**Table E6:** HMFAs were associated with atopy-implicated HMM traits.

| Estimate | P values[a] | $P_{Bonferroni}$[b] | 2.50% | 97.50% | HMM traits | HMFAs |
|---|---|---|---|---|---|---|
| **HMM network clusters** | | | | | | |
| -0.005 | 6.94E-04 | 2.08E-03 | -0.008 | -0.002 | *Green* | 24:0 |
| -0.004 | 7.84E-04 | 2.35E-03 | -0.006 | -0.002 | *Green* | 18:1c11 |
| -0.004 | 0.01 | 0.03 | -0.006 | -0.001 | *Green* | CLA |
| 0.003 | 0.03 | N/A | 0.0003 | 0.005 | *Green* | n-6/n-3 PUFAs |
| -0.003 | 0.01 | 0.03 | -0.005 | -0.001 | *Yellow* | 18:1c11 |
| 0.002 | 0.04 | N/A | 0.000 | 0.005 | *Yellow* | n-6/n-3 PUFAs |
| **CLR-transformed ASV abundance** | | | | | | |
| -0.25 | 1.94E-04 | 5.81E-04 | -0.38 | -0.12 | *Abiotrophia sp.* | 20:4n3 |
| -0.24 | 2.90E-04 | 8.71E-04 | -0.37 | -0.11 | *Abiotrophia sp.* | 20:5n3 |
| 0.24 | 1.80E-03 | N/A | 0.09 | 0.39 | *Granulicatella sp.* | n-6/n-3 PUFAs |
| -0.23 | 3.49E-03 | N/A | -0.38 | -0.08 | *Granulicatella sp.* | Total n-3 |
| -0.28 | 1.43E-03 | 4.30E-03 | -0.45 | -0.11 | *Granulicatella sp.* | CLA |
| -0.22 | 1.77E-03 | 5.32E-03 | -0.36 | -0.08 | *Abiotrophia sp.* | 22:6n3 |
| -0.23 | 2.56E-03 | 7.69E-03 | -0.38 | -0.08 | *Granulicatella sp.* | 20:5n3 |
| -0.20 | 3.08E-03 | 9.25E-03 | -0.34 | -0.07 | *Abiotrophia sp.* | 20:4n6 |
| -0.23 | 5.30E-03 | 0.02 | -0.39 | -0.07 | *Granulicatella sp.* | 22:6n3 |
| -0.21 | 6.68E-03 | 0.02 | -0.37 | -0.06 | *Granulicatella sp.* | 20:4n6 |
| -0.20 | 8.72E-03 | 0.03 | -0.35 | -0.05 | *Granulicatella sp.* | 20:4n3 |
| 0.15 | 0.03 | N/A | 0.02 | 0.28 | *Abiotrophia sp.* | n-6/n-3 PUFAs |
| -0.24 | 0.01 | 0.03 | -0.43 | -0.06 | *Haemophilus sp. 3* | 18:1c11 |
| -0.19 | 0.01 | 0.03 | -0.34 | -0.04 | *Granulicatella sp.* | 18:1c11 |
| -0.19 | 0.01 | 0.04 | -0.34 | -0.04 | *Granulicatella sp.* | 18:3n3 |
| 0.15 | 0.04 | N/A | 0.01 | 0.29 | *Abiotrophia sp.* | ARA/DHA |
| 0.19 | 0.05 | N/A | 0.00 | 0.38 | *Haemophilus sp. 3* | n6/n-3 PUFAs |
| 0.18 | 0.02 | 0.05 | 0.03 | 0.32 | *Granulicatella sp.* | 14:0 |
| **CLR-transformed MetaCyc Pathway abundance** | | | | | | |
| 0.14 | 1.05E-04 | 3.14E-04 | 0.07 | 0.21 | L-ornithine biosynthesis (GLUTORN-PWY) | 18:1c11 |
| 0.14 | 1.70E-04 | 5.09E-04 | 0.07 | 0.21 | L-histidine biosynthesis (HISTSYN-PWY) | 20:4n3 |

| | | | | | | |
|---|---|---|---|---|---|---|
| 0.12 | 1.33E-03 | 3.99E-03 | 0.05 | 0.19 | L-histidine biosynthesis (HISTSYN-PWY) | 20:5n3 |
| 0.10 | 1.59E-03 | 4.76E-03 | 0.04 | 0.17 | preQ0 biosynthesis (PWY-6703) | 18:1c11 |
| 0.18 | 1.61E-03 | 4.84E-03 | 0.07 | 0.29 | L-ornithine biosynthesis (GLUTORN-PWY) | 22:4n6 |
| 0.16 | 2.16E-03 | 6.48E-03 | 0.06 | 0.26 | preQ0 biosynthesis (PWY-6703) | 22:4n6 |
| -0.08 | 3.12E-03 | 0.01 | -0.13 | -0.03 | Superpathway of heme biosynthesis from glutamate (PWY-5918) | 10:0 |
| 0.12 | 4.26E-03 | 0.01 | 0.04 | 0.20 | Superpathway of heme biosynthesis from glutamate (PWY-5918) | 22:4n6 |
| 0.07 | 5.84E-03 | 0.02 | 0.02 | 0.12 | Partial TCA cycle (obligate autotrophs) (PWY-5913) | 20:5n3 |
| 0.09 | 0.01 | 0.03 | 0.02 | 0.17 | L-ornithine biosynthesis (GLUTORN-PWY) | 20:4n3 |
| -0.05 | 0.04 | N/A | -0.09 | -0.003 | Mixed acid fermentation (FERMENTATION-PWY) | Total saturated FA |
| 0.07 | 0.01 | 0.04 | 0.01 | 0.12 | TCA cycle VI (obligate autotrophs) (PWY-5913) | 22:6n3 |
| -0.08 | 0.02 | 0.05 | -0.15 | -0.02 | preQ0 biosynthesis (PWY-6703) | 10:0 |
| 0.06 | 0.02 | 0.05 | 0.01 | 0.11 | TCA cycle VI (obligate autotrophs) (PWY-5913) | 20:4n3 |
| 0.08 | 0.02 | 0.05 | 0.02 | 0.15 | preQ0 biosynthesis (PWY-6703) | 20:4n3 |
| -0.05 | 0.02 | 0.05 | -0.10 | -0.01 | Mixed acid fermentation (FERMENTATION-PWY) | 14:0 |
| -0.07 | 0.05 | N/A | -0.14 | 0.001 | L-histidine biosynthesis (HISTSYN-PWY) | n-6/n-3 PUFAs |

[a]Unadjusted P values were reported for ratios and total HMFAs.
[b]P values for 28 individual HMFAs were adjusted using Bonferroni correction for three clusters of correlated HMFAs.

**Table E7:** HMOs were associated with atopy-implicated HMM traits.

| Estimate | P values[a] | $P_{Bonferroni}$[b] | 2.50% | 97.50% | HMM traits | HMOs |
|---|---|---|---|---|---|---|
| | | | | | **HMM network clusters** | |
| -0.003 | 0.004 | 0.027 | -0.006 | -0.001 | *Green* | LNH |
| 0.003 | 0.008 | 0.046 | 0.003 | 0.001 | *Green* | 3FL |
| | | | | | **CLR-transformed ASV abundance** | |
| -0.33 | 8.76E-07 | 5.25E-06 | -0.46 | -0.2 | *Abiotrophia sp.* | DSLNH |
| -0.32 | 1.80E-06 | 1.08E-05 | -0.44 | -0.19 | *Abiotrophia sp.* | 6'SL |
| -0.24 | 0.0003 | 0.002 | -0.37 | -0.11 | *Abiotrophia sp.* | LNH |
| -0.24 | 0.002 | 0.01 | -0.39 | -0.09 | *Granulicatella sp.* | 6'SL |
| -0.24 | 0.002 | 0.01 | -0.39 | -0.09 | *Granulicatella sp.* | DSLNH |
| -0.23 | 0.002 | 0.01 | -0.38 | -0.09 | *Abiotrophia sp.* | LSTc |
| -0.19 | 0.006 | 0.04 | -0.32 | -0.05 | *Abiotrophia sp.* | FLNH |

[a]Unadjusted P values were reported for ratios and total HMOs.

[b]P values for 19 individual HMOs were adjusted using Bonferroni correction for six clusters of correlated HMOs.

**Table E8:** weighted gene co-expression network analysis (WGCNA) clustered 179 individual milk ASVs, 19 individual HMOs, and 28 HMFAs into seven multi-component network modules encompassing 226 milk features: 10 in multi-black, 49 in multi-blue, 39 in multi-brown, 32 in multi-purple, 15 in multi-red, 32 in multi-orange, and 49 in multi-turquoise.

| **Multi-black (n=10)** | **Multi-blue (n=49)** | **Multi-brown (n=39)** | **Multi-purple (n=32)** | **Multi-red (n=15)** | **Multi-orange (n=32)** | **Multi-turquoise (n=49)** |
|---|---|---|---|---|---|---|
| ***Staphylococcus sp.1**** | ***Abiotrophia sp.**** | **20:5n3*** | ***Streptococcus sanguinis**** | **15:0*** | ***Blastococcus sp.3**** | ***Stenotrophomonas maltophilia**** |
| *Anaerococcus sp.10* | *Actinomyces sp.1* | *Lactobacillus sp.23* | *Chryseobacterium sp.11* | FDSLNH | *Bradyrhizobium elkanii* | *Acinetobacter johnsonii* |
| *Peptoniphilus sp.1* | *Atopobium sp.* | *Rothia sp.3* | *Scardovia wiggsiae* | LNFP2 | *Comamonas aquatica* | *Squalius pyrenaicus* |
| *Corynebacterium sp.2* | *Prevotella sp.56* | 2'FL | *Veillonella sp.5* | LNFP3 | *Thermomonas sp.2* | *Aeromonas sp.* |
| *Corynebacterium striatum* | *Lachnoanaerobaculum sp.* | 6'SL | *Kocuria turfanensis* | LNT | *Methylobacterium-Methylorubrum sp.5* | *Lactobacillus sp.5* |
| *Corynebacterium sp.9* | *Lachnoanaerobaculum sp.1* | DFLNH | *Schlegelella aquatica* | LSTb | *proteobacterium symbiont* | *Pseudomonas sp.2* |
| *Corynebacterium sp.3* | *Veillonella sp.6* | DSLNH | *[Ruminococcus] gnavus sp.* | 10:0 | *Corynebacterium sp.16* | *Pseudomonas oryzihabitans* |
| *Finegoldia sp.* | *Prevotella salivae* | FLNH | *Romboutsia sp.1* | 12:0 | *Enhydrobacter sp.* | *Lactobacillus sp.8* |
| *Anaerococcus sp.7* | *Veillonella atypica* | LNFP1 | *Anaerococcus sp.1* | 14:0 | *Sphingobium sp.2* | *Rothia kristinae* |
| *Staphylococcus haemolyticus 2* | *Streptococcus sp.5* | LNH | *Anaerococcus sp.2* | 14:1n9 | *Aerococcus sp.* | *Novosphingobium sp.5* |
| | *Streptococcus sp.* | LNnT | *Thermus sp.* | DSLNT | *Psychrobacter immobilis* | *Unclassified Rhizobiaceae sp.3* |
| | *Veillonella sp.28* | LSTc | *Corynebacterium amycolatum* | 16:0 | *Massilia sp.1* | *Delftia sp.* |
| | 3FL | TVA | *Psychrobacter sanguinis* | 16:1n9 | *Auricoccus-Abyssicoccus sp.* | *Acinetobacter sp.1* |
| | 3'SL | 18:1n9 | *Abiotrophia sp.2* | 17:0 | *Brachybacterium sp.1* | *Pseudomonas sp.3* |
| | DFLac | 18:1C11 | *Veillonella sp.30* | 18:0 | *Skermanella sp.1* | *Pantoea sp.1* |
| | DFLNT | 18:2n6 | *Unclassified Neisseriaceae sp.* | | *Paracoccus sp.16* | *Enterobacter sp.3* |

| | | | | | | |
|---|---|---|---|---|---|---|
| | *Rothia sp.1* | 20:0 | *Corynebacterium sp.26* | | *Planococcus sp.* | *Sphingomonas sp.45* |
| | *Actinomyces sp.* | 18:3n6 | *Neisseria sp.4* | | *Caulobacter vibrioides* | *Chryseobacterium sp.* |
| | *Fusobacterium sp.2* | 18:3n3 | *Rothia sp.6* | | *Noviherbaspirillum sp.5* | *Acinetobacter sp.9* |
| | *Porphyromonas sp.6* | 20:2n6 | *Actinomyces sp.6* | | *Acinetobacter sp.* | *Methylorubrum rhodesianum* |
| | *Prevotella sp.37* | 20:3n6 | *Corynebacterium sp.* | | *Massilia sp.3* | *Exiguobacterium sp.1* |
| | *Alloprevotella sp.3* | 20:4n6 | *Neisseria sp.3* | | *Pseudomonas monteilii* | *Stenotrophomonas sp.* |
| | *Rothia sp.* | 20:4n3 | *Lautropia sp.* | | *Cloacibacterium sp.* | *Altererythrobacter sp.* |
| | *Bergeyella sp.3* | CLA | *Corynebacterium sp.1* | | *Bosea sp.1* | *Acinetobacter sp.23* |
| | *Moraxella sp.* | 24:0 | *Lawsonella sp.* | | *Pseudarthrobacter polychromogenes* | *Pantoea septica* |
| | *Dolosigranulum sp.* | 24:1n9 | *Anaerococcus sp.5* | | *Shewanella sp.1* | *Pseudomonas hunanensis* |
| | *Veillonella sp.10* | 22:4n6 | *Peptoniphilus sp.* | | *Pseudoalteromonas sp.1* | *Pantoea sp.* |
| | *Porphyromonas sp.14* | 22:5n6 | *Prevotella sp.6* | | *Bacillus sp.2* | *Pseudomonas sp.11* |
| | *Bergeyella sp.1* | 22:5n3 | *Fenollaria sp.* | | *Dietzia sp.2* | *Pseudoxanthomonas sp.5* |
| | *Haemophilus sp.15* | 22:6n3 | *Bifidobacterium catenulatum* | | *Bacillus niabensis* | *Chryseobacterium sp.5* |
| | *Gemella sp.* | *Rothia sp.4* | *Lactococcus lactis* | | *Paracoccus marcusii* | *Bacillus sp.23* |
| | *Granulicatella sp.* | *Sphingomonas sp.52* | *Brochothrix thermosphacta* | | *Micrococcus luteus 1* | *Rahnella sp.1* |
| | *Porphyromonas sp.11* | *Bifidobacterium longum 1* | | | | *Stenotrophomonas maltophilia 1* |
| | *Haemophilus sp.3* | *Escherichia coli* | | | | *Pseudomonas aeruginosa* |
| | *Prevotella sp.32* | *Klebsiella sp.4* | | | | *Klebsiella oxytoca* |

| | | | | | | |
|---|---|---|---|---|---|---|
| | *Prevotella sp.3* | *Lactobacillus fermentum* | | | | *Enterobacter cloacae* |
| | *Prevotella sp.38* | *Roseomonas gilardii* | | | | *Rahnella sp.* |
| | *Bergeyella sp.2* | *Bacillus subtilis* | | | | *Sphingomonas sp.26* |
| | *Megasphaera micronuciformis* | *Kocuria rhizophila* | | | | *Agrobacterium radiobacter* |
| | *Actinomyces sp.3* | | | | | *Pseudomonas viridiflava* |
| | *Campylobacter sp.2* | | | | | *Enterococcus faecalis 1* |
| | *Rothia sp.2* | | | | | *Brevundimonas sp.* |
| | *Veillonella sp.7* | | | | | *Stenotrophomonas sp.1* |
| | *Porphyromonas sp.10* | | | | | *Brevundimonas sp.6* |
| | *Leptotrichia sp.6* | | | | | *Acinetobacter baumannii* |
| | *Streptococcus salivarius* | | | | | *Sphingobacterium multivorum* |
| | *Streptococcus sp.2* | | | | | *Acinetobacter ursingii* |
| | *Corynebacterium propinquum* | | | | | *Pseudomonas sp.* |
| | *Neisseria perflava* | | | | | *Acinetobacter sp.40* |

*Hub milk features with the highest connectivity to other features within each module.

### Supplementary Figures

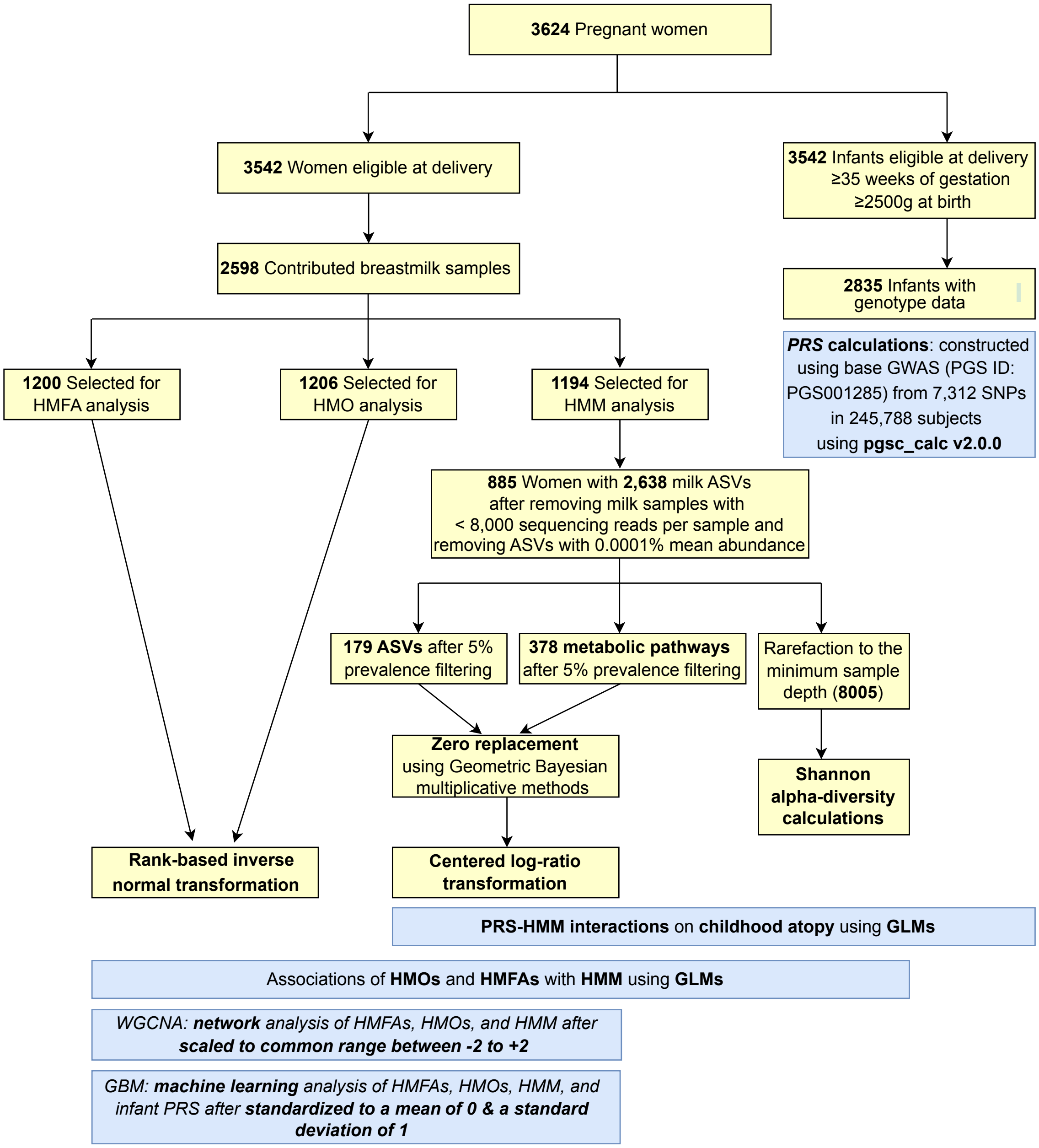

**Figure E1: Study flowchart of maternal milk sample and infant genomic analyses.**

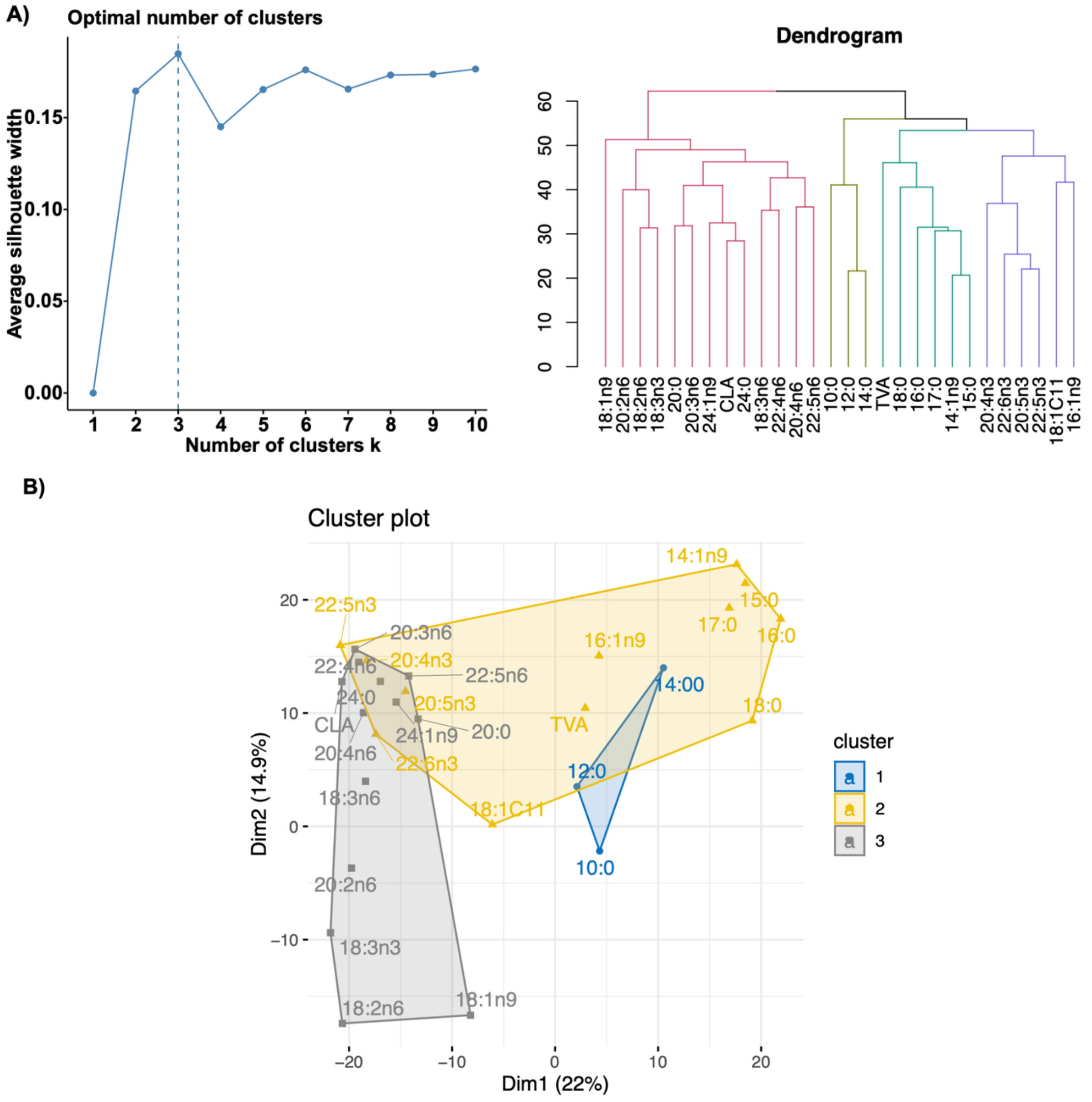

**Figure E2: Clustering of 28 individual HMFA profiles among 1,200 lactating mothers, related to Methods.** (**A**) Silhouette plots for determining the optimal number of clusters. The left panel shows the silhouette method applied to k-means clustering. (**B**) A dendrogram from hierarchical clustering grouped 28 individual HMFAs into three clusters based on Euclidean distance. (**C**) Cluster plot of 28 individual HMFAs projected onto the first two principal components, separating features into three clusters. Capric acid (10:0), lauric acid (12:0), myristic acid (14:0), physeteric acid (14:1n9), pentadecylic acid (15:0), palmitic acid (16:0), 7-hexadecenoic acid (16:1n9), heptadecanoic acid (17:0), stearic acid (18:0), trans-vaccenic acid (TVA; 18:1t11), oleic acid (18:1n9), cis-vaccenic acid (18:1c11), linoleic acid (LA; 18:2n6), arachidic acid (20:0), γ-linolenic acid (GLA; 18:3n6), α-linolenic acid (ALA; 18:3n3), eicosadienoic acid (20:2n6), dihomo-γ-linolenic acid (DGLA; 20:3n6), arachidonic acid (ARA; 20:4n6), conjugated linoleic acid (CLA; a family of linoleic acid isomers), eicosapentaenoic acid (EPA; 20:5n3), nervonic acid (24:1n9), docosahexaenoic acid (DHA; 22:6n3), lignoceric acid (24:0), osbond acid (22:5n6), docosapentaenoic acid (DPA; 22:5n3), eicosatetraenoic acid (20:4n3), and adrenic acid (22:4n6).

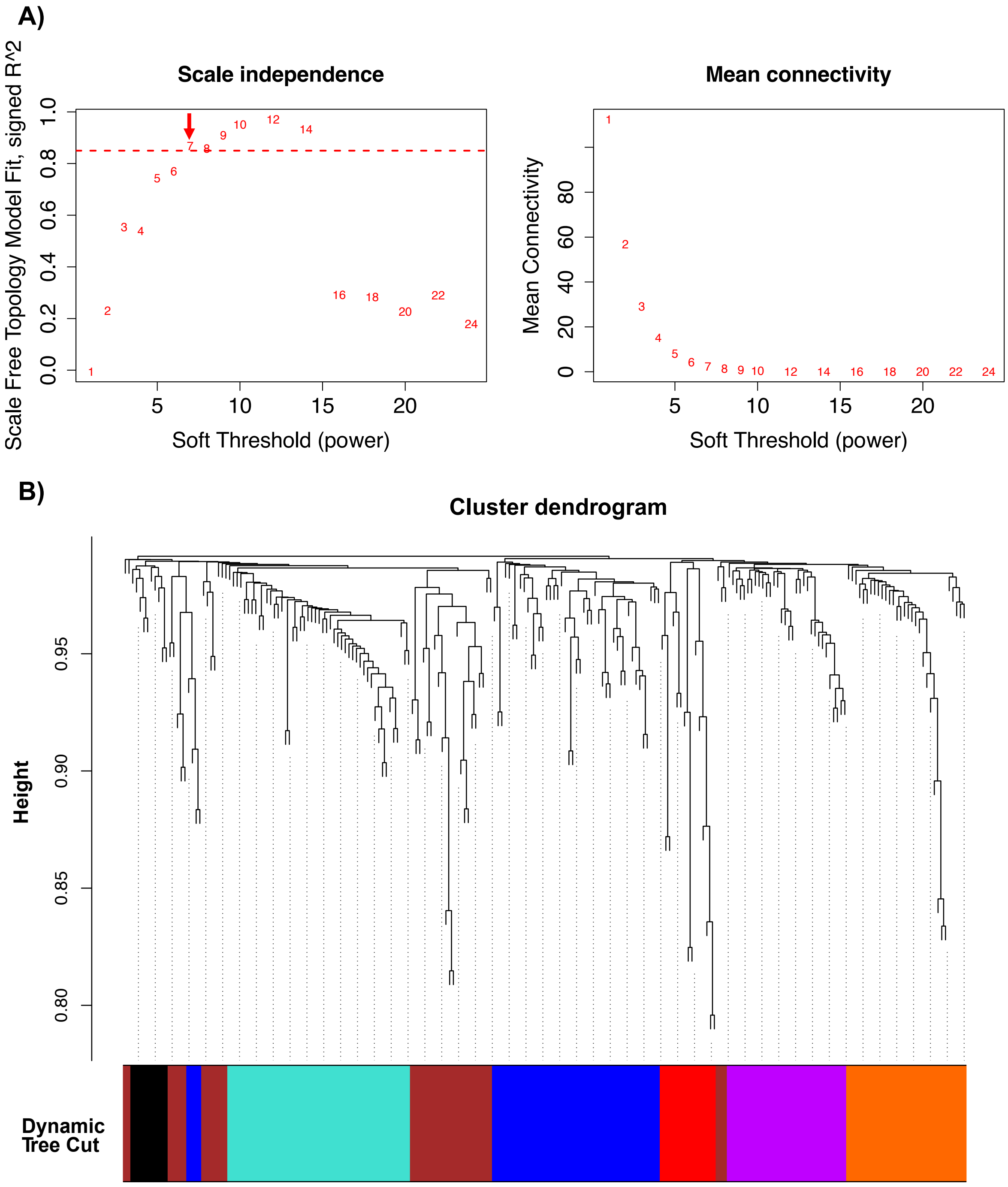

**Figure E3: WGCNA clustered 179 amplicon sequence variants (ASVs), 19 human milk oligosaccharides (HMOs), and 28 human milk fatty acids (HMFAs) into seven co-occurring network modules.** (**A**) Effects of candidate soft-thresholding powers on scale-free topology fit (left) and mean node connectivity (right). At β = 7, the co-abundance network modules of human milk components showed 86.8% similarity to a scale-free topology. At this β value, mean node connectivity also approximated a distribution typical of scale-free topology. The selected optimal β is marked with a red arrow in the left panel. (**B**) A dendrogram showing 179 ASV, 19 HMO, and 28 HMFA features clustered into seven network modules, as indicated by colours, using hierarchical clustering based on the topological overlap matrix (TOM).

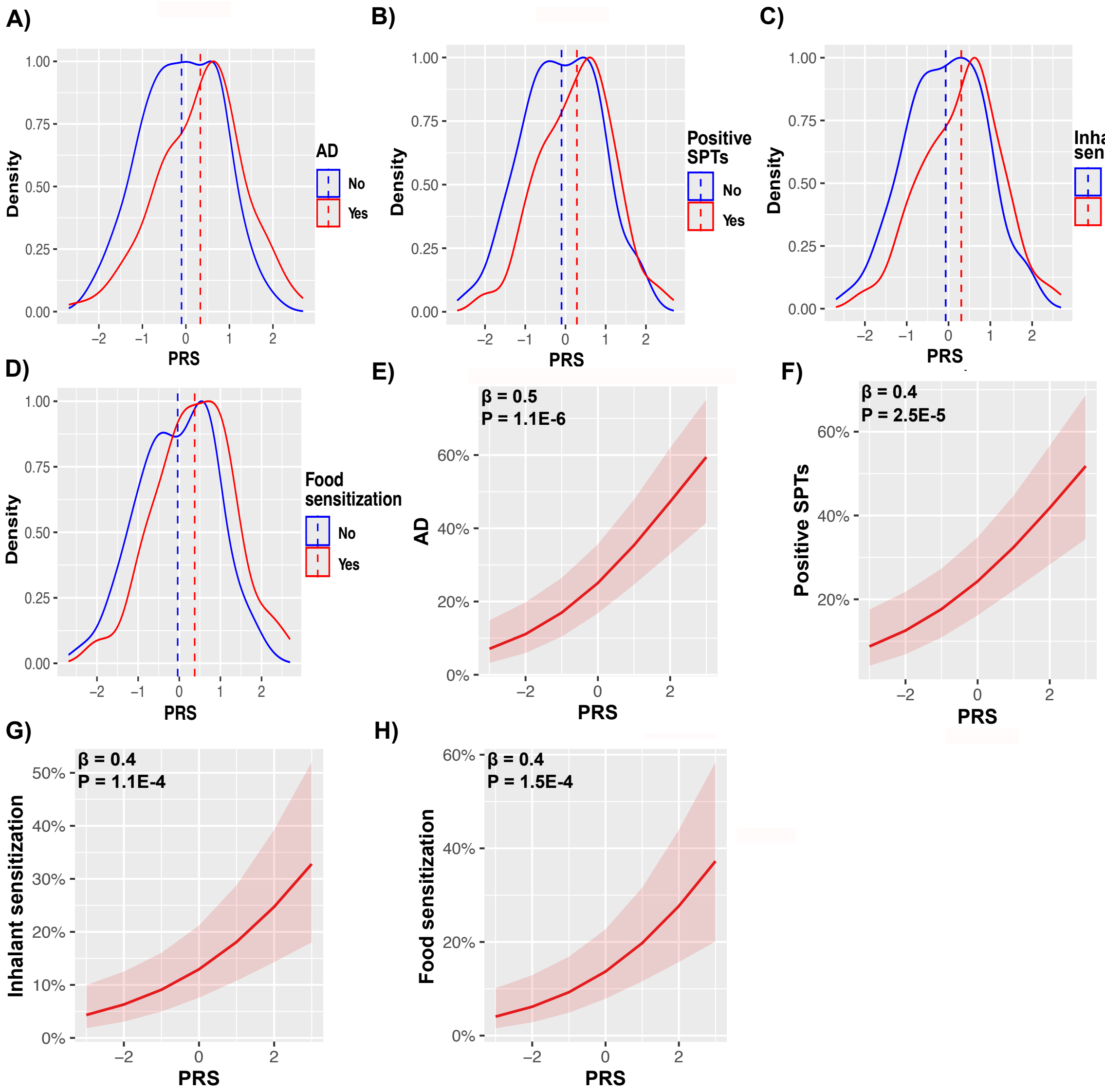

**Figure E4: Polygenic risk scores (PRS) were associated with childhood atopy at ages 1-5 years in the CHILD dataset.** Density plots show the PRS distribution among children with and without AD (**A**), food/inhalant sensitization (i.e., positive SPTs) (**B**), and allergic sensitization to food (**C**) and inhalant (**D**) allergens. Increased PRS was significantly associated with increased prevalence of childhood AD (**E**), food/inhalant sensitization (i.e., positive SPTs) (**F**), and allergic sensitization to food (**G**) and inhalant (**H**) allergens. Shaded areas represent the corresponding standard errors (SE).
